# Atrial Fibrillation Genetic Risk Differentiates Cardioembolic Stroke from other Stroke Subtypes

**DOI:** 10.1101/239269

**Authors:** Sara L. Pulit, Lu-Chen Weng, Patrick F McArdle, Ludovic Trinquart, Seung Hoan Choi, Braxton D. Mitchell, Jonathan Rosand, Paul I W de Bakker, Emelia J Benjamin, Patrick T Ellinor, Steven J Kittner, Steven A Lubitz, Christopher D Anderson, on behalf of the Atrial Fibrillation Genetics Consortium and the International Stroke Genetics Consortium, Ingrid E. Christophersen, Michiel Rienstra, Carolina Roselli, Xiaoyan Yin, Bastiaan Geelhoed, John Barnard, Honghuang Lin, Dan E. Arking, Albert V. Smith, Christine M. Albert, Mark Chaffin, Nathan R. Tucker, Molong Li, Derek Klarin, Nathan A Bihlmeyer, Siew-Kee Low, Peter E. Weeke, Martina Müller-Nurasyid, J. Gustav Smith, Jennifer A. Brody, Maartje N. Niemeijer, Marcus Dörr, Stella Trompet, Jennifer Huffman, Stefan Gustafsson, Claudia Schurmann, Marcus E. Kleber, Leo-Pekka Lyytikäinen, Ilkka Seppälä, Rainer Malik, Andrea R. V. R. Horimoto, Marco Perez, Juha Sinisalo, Stefanie Aeschbacher, Sébastien Thériault, Jie Yao, Farid Radmanesh, Stefan Weiss, Alexander Teumer, Seung Hoan Choi, Lu-Chen Weng, Sebastian Clauss, Rajat Deo, Daniel J. Rader, Svati Shah, Joylene E. Siland, Michiaki Kubo, Jonathan D. Smith, David R. Van Wagoner, Joshua C. Bis, Siegfried Perz, Bruce M. Psaty, Paul M. Ridker, Jared W. Magnani, Tamara B. Harris, Lenore J. Launer, M. Benjamin Shoemaker, Sandosh Padmanabhan, Jeffrey Haessler, Traci M. Bartz, Melanie Waldenberger, Peter Lichtner, Marina Arendt, Jose E. Krieger, Mika Kähönen, Lorenz Risch, Alfredo J. Mansur, Annette Peters, Blair H. Smith, Lars Lind, Stuart A. Scott, Yingchang Lu, Erwin B. Bottinger, Jussi Hernesniemi, Cecilia M. Lindgren, Jorge A Wong, Jie Huang, Markku Eskola, Andrew P. Morris, Ian Ford, Alex P. Reiner, Graciela Delgado, Lin Y. Chen, Yii-Der Ida Chen, Roopinder K. Sandhu, Man Li, Eric Boerwinkle, Lewin Eisele, Lars Lannfelt, Natalia Rost, arju Orho-Melander, Anders Hamsten, Jan Heeringa, Joshua C. Denny, Jennifer Kriebel, Dawood Darbar, Christopher Newton-Cheh, Christian Shaffer, Peter W. Macfarlane, Stefanie Heilmann, Peter Almgren, Paul L. Huang, Nona Sotoodehnia, Elsayed Z. Soliman, Andre G. Uitterlinden, Albert Hofman, Oscar H. Franco, Uwe Völker, Karl-Heinz Jöckel, Moritz F. Sinner, Henry J. Lin, Xiuqing Guo, Martin Dichgans, Erik Ingelsson, Charles Kooperberg, Olle Melander, Ruth J. F. Loos, Jari Laurikka, David Conen, Jonathan Rosand, Pim van der Harst, Marja-Liisa Lokki, Sekar Kathiresan, Alexandre Pereira, J. Wouter Jukema, Caroline Hayward, Jerome I. Rotter, Winfried März, Terho Lehtimäki, Bruno H. Stricker, Mina K. Chung, Stephan B. Felix, Vilmundur Gudnason, Alvaro Alonso, Dan M. Roden, Albert Sun, Christopher D. Anderson, Stefan Kääb, Jemma C. Hopewell, Stephanie Debette, Ganesh Chauhan, Qiong Yang, Bradford B. Worrall, Guillaume Paré, Yoichiro Kamatani, Yanick P. Hagemeijer, Niek Verweij, Kent D. Taylor, Archie Campbell, Patrik K. Magnusson, David Porteous, Lynne J. Hocking, Efthymia Vlachopoulou, Nancy L. Pedersen, Kjell Nikus, Daniel I. Chasman, Susan R. Heckbert, Emelia J. Benjamin, Toshihiro Tanaka, Kathryn L. Lunetta, Steven A. Lubitz, Patrick T. Ellinor, Sylvia Smoller, John Sorkin, Xingwu Wang, Magdy Selim, Aleksandra Pikula, Philip Wolf, Stephanie Debette, Sudha Seshadri, Paul de Bakker, Sara L. Pulit, Daniel Chasman, Kathryn Rexrode, Ida Chen, Jerome Rotter, May Luke, Michelle Sale, Tsong-Hai Lee, Ku-Chou Chang, Mitchell Elkind, Larry Goldstein, Michael Luke James, Monique Breteler, Chris O’Donnell, Didier Leys, Cara Carty, Chelsea Kidwell, Jes Olesen, Pankaj Sharma, Stephen Rich, Turgot Tatlisumak, Olli Happola, Philippe Bijlenga, Carolina Soriano, Eva Giralt, Jaume Roquer, Jordi Jimenez-Conde, Ioana Cotlarcius, John Hardy, Michal Korostynski, Giorgio Boncoraglio, Elena Ballabio, Eugenio Parati, Adamski Mateusz, Andrzej Urbanik, Tomasz Dziedzic, Jeremiasz Jagiella, Jerzy Gasowski, Marcin Wnuk, Rafael Olszanecki, Joanna Pera, Agnieszka Slowik, Karol Jozef Juchniewicz, Christopher Levi, Paul Nyquist, Iscia Cendes, Norberto Cabral, Paulo Franca, Anderson Goncalves, Lina Keller, Milita Crisby, Konstantinos Kostulas, Robin Lemmens, Kourosh Ahmadi, Christian Opherk, Marco Duering, Martin Dichgans, Rainer Malik, Mariya Gonik, Julie Staals, Olle Melander, Philippe Burri, Ariane Sadr-Nabavi, Javier Romero, Alessandro Biffi, Chris Anderson, Guido Falcone, Bart Brouwers, Jonathan Rosand, Natalia Rost, Rose Du, Christina Kourkoulis, Thomas Battey, Steven Lubitz, Bertram Mueller-Myhsok, James Meschia, Thomas Brott, Guillaume Pare, Alexander Pichler, Christian Enzinger, Helena Schmidt, Reinhold Schmidt, Stephan Seiler, Susan Blanton, Yoshiji Yamada, Anna Bersano, Tatjana Rundek, Ralph Sacco, Yu-Feng Yvonne Chan, Andreas Gschwendtner, Zhen Deng, Taura Barr, Katrina Gwinn, Roderick Corriveau, Andrew Singleton, Salina Waddy, Lenore Launer, Christopher Chen, Kim En Le, Wei Ling Lee, Eng King Tan, Akintomi Olugbodi, Peter Rothwell, Sabrina Schilling, Vincent Mok, Elena Lebedeva, Christina Jern, Katarina Jood, Sandra Olsson, Helen Kim, Chaeyoung Lee, Laura Kilarski, Hugh Markus, Jennifer Peycke, Steve Bevan, Wayne Sheu, Hung Yi Chiou, Joseph Chern, Elias Giraldo, Muhammad Taqi, Vivek Jain, Olivia Lam, George Howard, Daniel Woo, Steven Kittner, Braxton Mitchell, John Cole, Jeff O’Connell, Dianna Milewicz, Kachikwu Illoh, Bradford Worrall, Colin Stine, Bartosz Karaszewski, David Werring, Reecha Sofat, June Smalley, Arne Lindgren, Bjorn Hansen, Bo Norrving, Gustav Smith, Juan Jose Martin, Vincent Thijs, Karin Klijn, Femke van’t Hof, Ale Algra, Mary Macleod, Rodney Perry, Donna Arnett, Alessandro Pezzini, Alessandro Padovani, Steve Cramer, Mark Fisher, Danish Saleheen, Joseph Broderick, Brett Kissela, Alex Doney, Sudlow Cathie, Kristiina Rannikmae, Scott Silliman, Caitrin McDonough, Matthew Walters, Annie Pedersen, Kazuma Nakagawa, Christy Chang, Mark Dobbins, Patrick McArdle, Yu-Ching Chang, Robert Brown, Devin Brown, Elizabeth Holliday, Raj Kalaria, Jane Maguire, Attia John, Martin Farrall, Anne-Katrin Giese, Myriam Fornage, Jennifer Majersik, Mary Cushman, Keith Keene, Siiri Bennett, David Tirschwell, Bruce Psaty, Alex Reiner, Will Longstreth, David Spence, Joan Montaner, Israel Fernandez-Cadenas, Carl Langefeld, Cheryl Bushnell, Laura Heitsch, Jin-Moo Lee, Kevin Sheth

## Abstract

**Objective:** We sought to assess whether genetic risk factors for atrial fibrillation can explain cardioembolic stroke risk.

**Methods:** We evaluated genetic correlations between a prior genetic study of AF and AF in the presence of cardioembolic stroke using genome-wide genotypes from the Stroke Genetics Network (N = 3,190 AF cases, 3,000 cardioembolic stroke cases, and 28,026 referents). We tested whether a previously-validated AF polygenic risk score (PRS) associated with cardioembolic and other stroke subtypes after accounting for AF clinical risk factors.

**Results:** We observed strong correlation between previously reported genetic risk for AF, AF in the presence of stroke, and cardioembolic stroke (Pearson’s r=0.77 and 0.76, respectively, across SNPs with p < 4.4 × 10^−4^ in the prior AF meta-analysis). An AF PRS, adjusted for clinical AF risk factors, was associated with cardioembolic stroke (odds ratio (OR) per standard deviation (sd) = 1.40, p = 1.45×10^−48^), explaining ∼20% of the heritable component of cardioembolic stroke risk. The AF PRS was also associated with stroke of undetermined cause (OR per sd = 1.07, p = 0.004), but no other primary stroke subtypes (all p > 0.1).

**Conclusions:** Genetic risk for AF is associated with cardioembolic stroke, independent of clinical risk factors. Studies are warranted to determine whether AF genetic risk can serve as a biomarker for strokes caused by AF.

## Introduction

Atrial fibrillation affects nearly 34 million individuals worldwide^1^ and is associated with a five-fold increased risk of ischemic stroke,^2^ a leading cause of death and disability.^3,4^ Atrial fibrillation promotes blood clot formation in the heart which can embolize distally, and is a leading cause of cardioembolism. Secondary prevention of cardioembolic stroke is directed at identifying atrial fibrillation as a potential cause, and initiating anticoagulation to prevent recurrences. Yet atrial fibrillation can remain occult even after extensive workup owing to the paroxysmal nature and fact that it can be asymptomatic. Since both atrial fibrillation and stroke are heritable, and since there is a compelling clinical need to determine whether stroke survivors have atrial fibrillation as an underlying cause, we sought to determine whether genetic risk of cardioembolic stroke can be approximated by measuring genetic susceptibility to atrial fibrillation.

Recent genome-wide association studies (GWAS) have demonstrated that both atrial fibrillation^5^ and ischemic stroke^6,7^ are complex disorders with polygenic architectures. The top loci for cardioembolic stroke, on chromosome 4q25 upstream of *PITX2* and on 16q22 near *ZFHX3*, are both leading risk loci for atrial fibrillation.^8–10^ Despite overlap in top risk loci, the genetic susceptibility to both atrial fibrillation and cardioembolic stroke is likely to involve the aggregate contributions of hundreds or thousands of loci, consistent with other polygenic conditions.^11^

To understand whether genetic risk for atrial fibrillation is an important and potentially useful determinant of overall cardioembolic stroke risk, we analyzed 13,390 ischemic stroke cases and 28,026 referents from the NINDS-Stroke Genetics Network (SiGN)^12^ with genome-wide genotyping data. First, we assessed whether stroke patients with atrial fibrillation have a genetic predisposition to the arrhythmia, leveraging additional GWAS data from the Atrial Fibrillation Genetics Consortium (AFGen). Second, we compared genetic risk factors for atrial fibrillation and stroke to ascertain the extent to which heritable risk of cardioembolic stroke is explained by genetic risk factors for atrial fibrillation.

## Methods

### The Stroke Genetics Network (SiGN)

The Stroke Genetics Network (SiGN) was established with the aim of performing the largest genome-wide association study (GWAS) of ischemic stroke to date. The study design has been described previously^12^ and is summarized in the **Supplementary Methods**. Briefly, subjects in SiGN were classified into stroke subtypes using the Causative Classification System (CCS), which subtypes cases through an automated, web-based system that accounts for clinical data, test results, and imaging information.^13,14^ Within CCS, there are two sub-categories: CCS causative, which does not allow for competing subtypes in a single sample; and CCS phenotypic, which does. Additionally, ∼74% of samples were subtyped using the TOAST subtyping system.^15^ After quality control (QC), the SiGN dataset comprised 16,851 ischemic stroke cases and 32,473 stroke-free controls (**Supplementary Methods** and **Supplementary Table 1**). Here, we analyze only the European-and African-ancestry samples (13,390 cases and 28,026 controls).

**Table 1.** Atrial fibrillation and stroke cases in SiGN. Of the 13,390 stroke cases available in the SiGN dataset, a total of 3,190 cases had atrial fibrillation or other suggestive diagnoses. While the majority of these cases were subtyped as having a cardioembolic stroke, a fraction was distributed among the other stroke subtypes. Samples can appear more than once per row (i.e., have more than one atrial fibrillation diagnosis), but totals represent the number of unique atrial fibrillation samples in each stroke subtype. There are no subjects with atrial fibrillation or equivalent subtyped as “cryptogenic/cardioembolic minor” because such a diagnosis would remove them from this category.

| Phenotype | Total | Ischemic stroke subtype |  |  |  |  |
| --- | --- | --- | --- | --- | --- | --- |
|  |  | Primary subtypes |  |  | Undetermined subtypes |  |
|  |  | Cardioembolic | Large artery atherosclerosis | Small artery occlusion | Incomplete/unclassified | Cryptogenic/cardioembolic minor |
| Atrial fibrillation | 1,751 | 1,495 | 63 | 32 | 151 | 0 |
| Paroxysmal atrial fibrillation | 1,315 | 1,088 | 52 | 23 | 138 | 0 |
| Left atrial thrombus | 48 | 37 | 3 | 3 | 4 | 0 |
| Sick sinus syndrome | 79 | 65 | 5 | 3 | 4 | 0 |
| Atrial Flutter | 106 | 90 | 4 | 2 | 10 | 0 |
| Total atrial fibrillation cases | 3,190 | 2,684 | 123 | 61 | 298 | 0 |
| No atrial fibrillation | -- | 316 | 2,262 | 2,201 | 1,982 | 2,294 |

### Standard Protocol Approvals, Registrations, and Patient Consents

All cohorts included in the SiGN dataset received approval from the cohort-specific ethical standards committee. Cohorts received written informed consent from all patients or guardians of patients participating in the study, where applicable. Details on sample collection have been described previously.^12^

### Identifying atrial fibrillation cases and controls

We defined atrial fibrillation in SiGN on the basis of five variables available in the CCS phenotyping system: (i) atrial fibrillation, (ii) paroxysmal atrial fibrillation, (iii) atrial flutter, (iv) sick sinus syndrome, and (v) atrial thrombus. This definition yielded 3,190 atrial fibrillation cases for analysis. We also defined a strict case set based on “atrial fibrillation” only (N = 1,751 cases) for sensitivity analyses (**Supplementary Methods** and **Supplementary Figure 1**).

**Figure 1.**
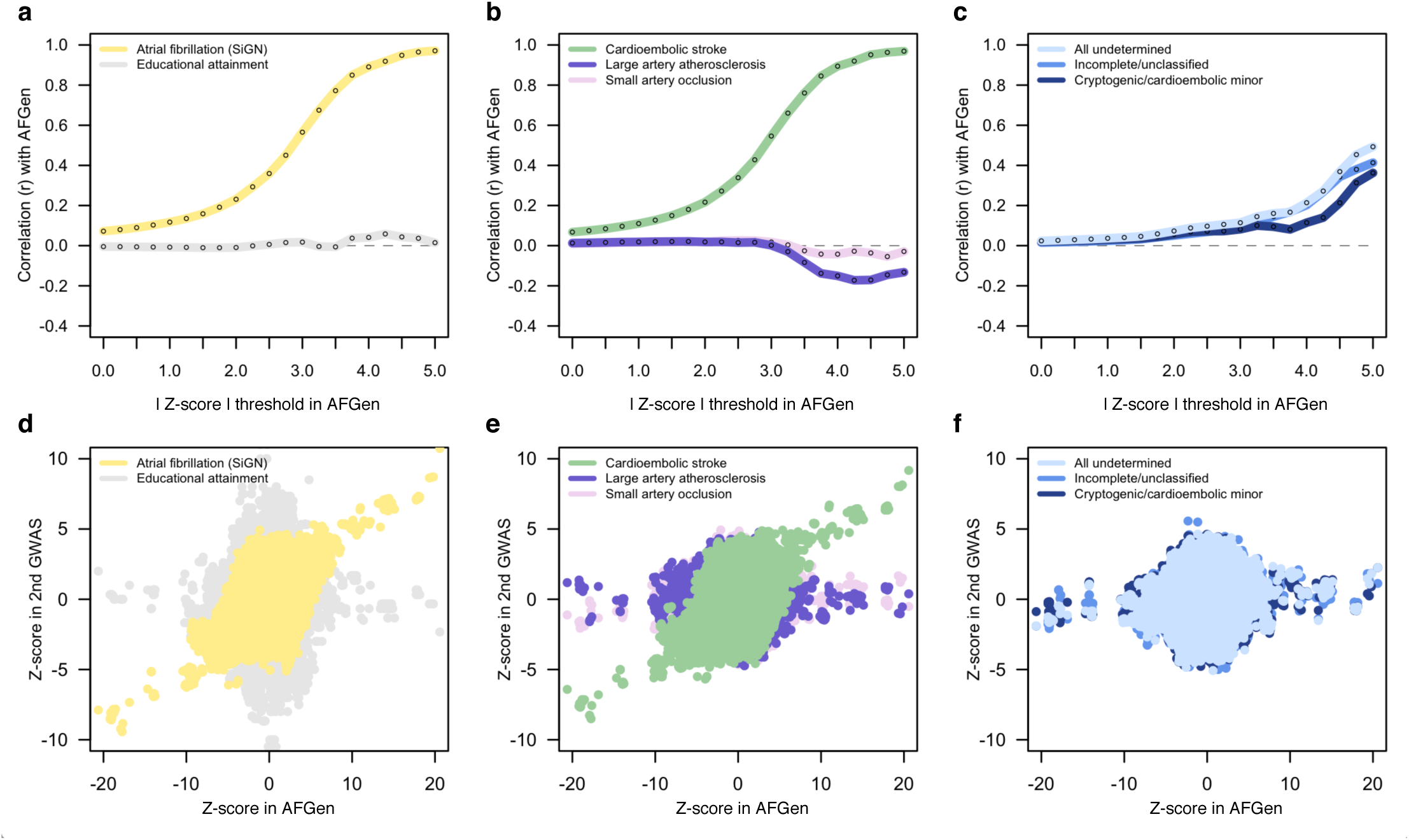
Genetic correlation between atrial fibrillation in the Atrial Fibrillation Genetics (AFGen) Consortium meta-analysis and atrial fibrillation and ischemic stroke subtypes analysed in SiGN. Pearson’s r correlation between SNP z-scores in the AFGen GWAS of atrial fibrillation and in GWAS of selected traits performed in the SiGN data. (a) GWAS of atrial fibrillation in AFGen and in SiGN correlate with increasing strength as SNP z-scores in AFGen increase. Correlation with educational attainment (performed separately, shown here as a null comparator) remains approximately zero across all z-score thresholds. (b) SNP effects in AFGen also correlate strongly with cardioembolic stroke in SiGN, but not with the other primary stroke subtypes. (c) Undetermined subtypes of stroke also show modest correlation to the genetic architecture of atrial fibrillation in AFGen. Panels d-f show genome-wide z-score distributions underlying correlations.

From the 28,026 controls, we established a set of 3,861 control individuals in whom atrial fibrillation was indicated as not present. For the remaining subjects, we assumed that individuals did not have atrial fibrillation since atrial fibrillation status for most control samples in SiGN is unknown.

### Genome-wide association testing of ischemic stroke subtypes and atrial fibrillation in SiGN

We merged genotype dosages together and kept SNPs with imputation quality > 0.8 and minor allele frequency (MAF) > 1% (**Supplementary Methods**). We performed association testing using a linear mixed model implemented in BOLT-LMM.^16^ We adjusted the model for the top ten principal components (PCs) and sex, in addition to the genetic relationship matrix (GRM; **Supplementary Methods**).^16^ We performed GWAS in atrial fibrillation and each of the stroke subtypes available in SiGN. Results were unadjusted for age, as adjusting for age in the atrial fibrillation GWAS gave results highly concordant with the age-unadjusted results (**Supplementary Results**).

### Heritability calculations

We calculated additive SNP-based heritability estimates for ischemic stroke, stroke subtypes, and atrial fibrillation using restricted maximum likelihood implemented in BOLT-REML (**Supplementary Methods**).^16^

### Genetic correlation between atrial fibrillation and ischemic stroke subtypes

We used summary-level data from a prior Atrial Fibrillation Genetics (AFGen) Consortium meta-analysis of atrial fibrillation^5^ to calculate a z-score for each SNP in that GWAS. Additionally, we calculated a z-score for each SNP from our SiGN GWAS of each stroke subtype and atrial fibrillation. As a null comparator, we downloaded SNP z-scores from a GWAS of educational attainment^17^ available through LDHub (http://ldsc.broadinstitute.org/, accessed 11-1-2017). We calculated Pearson’s r between z-scores from two traits to evaluate correlation (**Supplementary Methods** and **Supplementary Figure 3**).

### Constructing an atrial fibrillation polygenic risk score

To construct an atrial fibrillation polygenic risk score (PRS), we used SNPs from a previously-derived atrial fibrillation PRS (**Supplementary Methods**).^18^ Briefly, the PRS was derived from an atrial fibrillation GWAS of 17,931 cases and 115,142 controls.^5^ This PRS comprised 1,168 SNPs with p < 1 × 10^−4^ and LD pruned at an r^2^ threshold of 0.5.^18^ Of these 1,168 SNPs, we identified 934 SNPs in the SiGN dataset with imputation info > 0.8 and MAF > 1%. We used these 934 SNPs to construct the atrial fibrillation PRS in the SiGN dataset. Additional details on the PRS construction can be found in the **Supplementary Methods**.

### Testing an atrial fibrillation polygenic risk score in ischemic stroke subtypes

We tested for association between the atrial fibrillation PRS and stroke subtypes using logistic regression (**Supplementary Methods**). We included sex and the top 10 PCs as additional covariates. We optionally adjusted the association tests for age, diabetes mellitus, cardiovascular disease, smoking status (current smoker, former smoker, or never smoked), and hypertension.

We calculated the variance explained by the atrial fibrillation PRS in cardioembolic stroke by constructing a model in BOLT-REML that consisted of: (1) a variance component made up of SNPs for the GRM, and (2) a variance component made up of SNPs from the PRS (**Supplementary Methods**).

### Data availability

Code, supporting data, and downloadable supplemental tables are available here: https://github.com/UMCUGenetics/Afib-Stroke-Overlap. The **Supplementary Information** contains additional information regarding data access, methods, and links to summary-level data.

## Results

We began by testing our ability to rediscover known atrial fibrillation genetic associations in the SiGN dataset, assembled to study the genetics of ischemic stroke. We ran a genome-wide association study (GWAS) in SiGN using 3,190 cases with atrial fibrillation or paroxysmal atrial fibrillation, as well as other diagnoses suggestive of underlying atrial fibrillation^19,20^ (**Methods**, **Table 1** and **Supplementary Table 1**) and 28,026 controls (**Supplementary Figure 1**). We found the top-associated SNPs to be highly concordant with a prior GWAS of atrial fibrillation performed by the Atrial Fibrillation Genetics (AFGen) Consortium (**Supplementary Table 2**). Adjusting the GWAS for age did not substantially change our findings (r = 0.83 between SNP effects from the age-unadjusted and age-adjusted GWAS).

Extending our analysis beyond these top associations, we next assessed whether stroke patients with atrial fibrillation have a similar overall genetic predisposition to the arrhythmia as seen in the independent AFGen GWAS. Additionally, we assessed the overlap between genetic predisposition to atrial fibrillation and each stroke subtype, allowing for the known phenotypic concordance between cardioembolic stroke and atrial fibrillation (89.5% of cardioembolic stroke cases in SiGN also have atrial fibrillation, **Supplementary Table 1**). We performed a series of GWAS in the SiGN data for atrial fibrillation and each of the stroke subtypes using BOLT-LMM^16^ (**Methods**), and calculated the z-score (beta/standard error) of each SNP in each phenotype. We then used summary-level results available from the prior (independent) GWAS of atrial fibrillation^5^ (from AFGen) and calculated the z-score for each SNP in that dataset.

Measuring Pearson’s correlation (r) between AFGen z-scores and z-scores from the atrial fibrillation GWAS in SiGN, we found only a modest correlation (r =0.07 across ∼7.8M SNPs, **Figure 1**). However, when we iteratively subsetted the AFGen GWAS results by the (absolute values of) z-scores of the SNPs, we found that correlation with the atrial fibrillation GWAS in SiGN increased as the z-score threshold became more stringent. For example, for those ∼4.5M SNPs with |z| > 1 in AFGen, correlation with atrial fibrillation SNPs in SiGN was 0.12; for those ∼1.9M SNPs with |z| > 3.5 in AFGen, correlation with the SiGN atrial fibrillation GWAS rose to 0.77 (**Figure 1** and **Supplementary Table 3**). These correlations, calculated to include even modestly-associated SNPs, indicate that atrial fibrillation in AFGen and atrial fibrillation in stroke (SiGN) share a large proportion of genetic risk factors. Removing ±2Mb around the *PITX2* and *ZFHX3* loci only modestly impacted the correlation between AFGen and atrial fibrillation in SiGN (r = 0.63 for SNPs with |z| > 3.5; **Supplementary Figure 2** and **Supplementary Table 3**). Correlations between AFGen and cardioembolic stroke in SiGN were unsurprisingly highly similar to that of the results with atrial fibrillation in SiGN (r = 0.77 for AFGen SNPs with |z| > 3.5), likely due to the high concordance between the atrial fibrillation and cardioembolic stroke phenotypes (**Figure 1** and **Supplementary Figure 3**).

Continuing this analysis across the other stroke subtypes (large artery atherosclerosis, small artery occlusion, and undetermined stroke; **Figure 1**), we found near-zero correlation between AFGen and either large artery atherosclerosis or small artery occlusion (**Figure 1**) indicating no genetic overlap between the phenotypes. However, the correlation between atrial fibrillation and the undetermined stroke subtypes (a highly heterogeneous subset of cases^21,22^ that cannot be classified with standard subtyping systems^13,15^) increased steadily as we partitioned the AFGen data by z-score (all undetermined vs. AFGen r = 0.04 for AFGen SNPs with |z| > 1 and r = 0.16 for AFGen SNPs with |z| > 3.5; **Figure 1** and **Supplementary Table 3**), indicating that genome-wide, there is residual genetic correlation between atrial fibrillation and the undetermined stroke categories, some of which could represent causal atrial fibrillation stroke mechanisms in that subgroup. As an additional null comparator, we performed correlations between the AFGen results with z-scores derived from the latest GWAS of educational attainment^17^ and found that correlation remained at approximately zero regardless of the z-score threshold used (**Figure 1** and **Supplementary Table 3**).

To further understand the overlap between genetic risk factors for atrial fibrillation and cardioembolic stroke and to evaluate the degree to which cardioembolic stroke is comprised of risk factors beyond those for atrial fibrillation, we performed a restricted maximum likelihood analysis implemented in BOLT-REML^16^ to estimate SNP-based heritability of atrial fibrillation and cardioembolic stroke. Using phenotypes derived from the CCS subtyping algorithm^23^ (**Methods**), we estimated heritability of atrial fibrillation and cardioembolic stroke at 20.0% and 19.5%, respectively. These estimates are consistent with previous estimates in larger samples (**Supplementary Figure 4**),^24,25^ and the similar heritabilities suggest that cardioembolic stroke does not have a substantial heritable component beyond the primary atrial fibrillation risk factor. For comparison, we calculated heritability in the other stroke subtypes^15^ and found estimates to be similarly modest (range: 15.5% - 23.0%; **Supplementary Figures 4-6** and **Supplementary Table 4**).

Up to this point, our results indicated that atrial fibrillation in ischemic stroke is genetically similar to that discovered in prior genetic studies of atrial fibrillation alone, and that the bulk of the genetic risk for cardioembolic stroke appears attributable to atrial fibrillation genetic risk factors. Next, we sought to explicitly test what proportion of cardioembolic stroke risk could be explained by atrial fibrillation loci, independent of known clinical risk factors for atrial fibrillation. First, we identified SNPs from an atrial fibrillation polygenic risk score (PRS) independently derived from the AFGen GWAS^5^ (**Methods**). Of the 1,168 SNPs used to generate this pre-established PRS, we identified 934 in the SiGN dataset with imputation quality > 0.8 and minor allele frequency >1%. We computed the PRS per individual (**Methods**), weighting the imputed dosage of each risk allele by the effect of the SNP (i.e., the beta coefficient) as reported in AFGen^5^.

We tested the association of the atrial fibrillation PRS with cardioembolic stroke, using a logistic regression and adjusting for the top ten principal components and sex (**Methods**). As expected from our earlier results, we found the PRS to be strongly associated with cardioembolic stroke (odds ratio (OR) per 1 standard deviation (sd) of the PRS = 1.93 [95% confidence interval (CI): 1.34 - 1.44], p = 1.01 × 10^−65^; **Figure 2** and **Supplementary Table 5**), confirming the high genetic concordance of these phenotypes across SNPs which, individually, confer only a modest average association with atrial fibrillation. Next, we adjusted the association model for clinical covariates associated with atrial fibrillation including age, diabetes mellitus, cardiovascular disease, smoking, and hypertension.^26^ Using a (smaller) set of cases and controls with complete clinical risk factor information, we found that inclusion of these clinical risk factors in the model only modestly reduced the PRS signal in cardioembolic stroke (OR per 1 sd = 1.40 [95% CI: 1.34 - 1.47], p = 1.45 × 10^−48^; **Supplementary Tables 5**-**7**). These results indicate a strong relationship between atrial fibrillation genetic risk factors and cardioembolic stroke risk, independent of the clinical factors that associate with atrial fibrillation. Expanding the set of SNPs used to construct the PRS to the original 934 SNPs ±25kb, ±50kb, and ±100kb (**Methods**) revealed a persistently strong, though somewhat attenuated, association between the PRS and cardioembolic stroke (PRS including SNPs within 100kb, p = 4.47 × 10^−44^, **Supplementary Table 6**). None of the other stroke subtypes were significantly associated with the atrial fibrillation PRS (all p > 0.013, **Figure 2** and **Supplementary Figure 6**).

**Figure 2.**
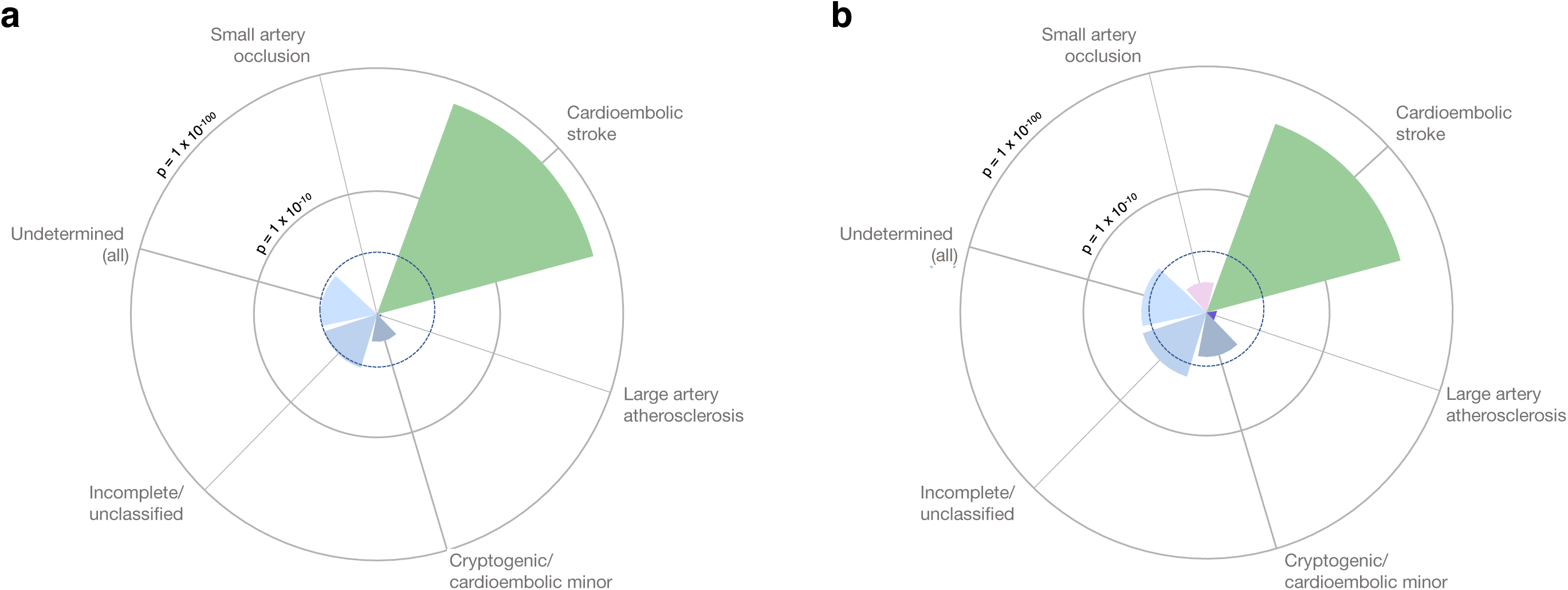
Association of atrial fibrillation polygenic risk score in ischemic stroke subtypes. We constructed an independent polygenic risk score (PRS) from atrial fibrillation-associated SNPs identified in the AFGen GWAS, and tested associations between this PRS and ischemic stroke subtypes using (a) all available referents (N = 28,026) and (b) referents without atrial fibrillation (N = 3,861). The PRS strongly associated with cardioembolic stroke in both sets of samples. In the atrial fibrillation-free set of controls (panel b) we observed association of the PRS (p < 5 × 10^−3^, after adjusting for five subtypes and two sets of referents; indicated by the dashed dark blue line) with incomplete/unclassified stroke as well.

Because atrial fibrillation status was missing for most controls in the SiGN dataset, we performed sensitivity analyses using only the 3,861 controls confirmed as having no atrial fibrillation. While reducing the set of controls to this refined group did not substantially change results for the primary stroke subtypes, we found the atrial fibrillation PRS was modestly associated (p < 5 × 10^−3^, after adjusting for five subtypes and two control groups) with the overall undetermined subtype (OR per 1 sd = 1.07 [95% CI: 1.02 - 1.13], p = 4.15 × 10^−3^) (**Figure 2** and **Supplementary Table 5**). Further examination of the two mutually exclusive subgroups of the undetermined group revealed that the PRS associated significantly with the incomplete/unclassified categorization (OR per 1 sd = 1.09 [95% CI: 1.03 - 1.16], p = 3.17 × 10^−3^) (**Figure 2**) but not with cryptogenic/cardioembolic minor (OR per 1 sd = 1.06 [95% CI: 1.00 - 1.13], p = 5.10 × 10^−2^). Correcting for clinical covariates only modestly changed the signal in the incomplete/unclassified phenotype (p = 9.7 × 10^−3^, **Figure 2**), supporting the robustness of the observed association, independent of clinical risk factors.

Lastly, we created a model in BOLT-LMM, fitting two genetic variance components: one component including SNPs for the genetic relationship matrix, and the second component including the original PRS SNPs from the atrial fibrillation PRS (including ±100kb around these SNPs, to include a sufficient number of markers to estimate variance explained). We found that the SNPs from the atrial fibrillation PRS explained 4.1% of the total (20.0%) heritability in atrial fibrillation. In evaluating variance explained in cardioembolic stroke, we found a nearly identical result: the component representing the atrial fibrillation risk score explained 4.5% (s.e. = 1.00%) of the total 19.5% genetic heritability in cardioembolic stroke. Thus, atrial fibrillation genetic risk accounts for 23.1%, or approximately one-fifth, of the total heritability of cardioembolic stroke.

## Discussion

Our results suggest that individuals with cardioembolic strokes have an enrichment for atrial fibrillation genetic risk, despite the fact that cardioembolic stroke often affects older adults with multiple clinical comorbidities^27^ that could increase risk for atrial fibrillation due to non-genetic factors. The fact that cardioembolic stroke and atrial fibrillation share a highly-similar genetic architecture extends our understanding of the morbid consequences of heritable forms of the arrhythmia. Furthermore, the observation that atrial fibrillation genetic risk was only associated with cardioembolic stroke, and (consistently) lacked association in large artery atherosclerosis or small artery occlusion,^28^ raises the possibility that atrial fibrillation genetic risk may be informative in the management of ischemic stroke survivors in whom the mechanism may be unclear.

The use of polygenic risk scores for complex traits has proved an efficient means of understanding how genetic predisposition to diseases can overlap. Given the onslaught of genotyping data available for common diseases, PRS’s can now be used to stratify patients by risk (e.g., in breast cancer^29,30^) or predict outcome (e.g., in neuropsychiatric disease^29^). More recently, PRS’s have been used to identify individuals in the general population with a four-fold risk for coronary disease,^31^ proposed for inclusion in clinical workups of individuals with early-onset coronary artery disease,^32^ and used to identify patients for whom lifestyle changes or statin intervention would be beneficial.^33,34^ While previous work has also shown an association between an atrial fibrillation PRS and cardioembolic stroke,^28^ we have extended this work to formally quantify the extent to which an atrial fibrillation PRS captures genetic risk for cardioembolic stroke. These findings lay the groundwork for future work that can potentially leverage this overlap to develop atrial fibrillation PRS’s that could be used to predict individuals at highest risk of cardioembolic stroke (to improve diagnostic resource allocation) or help distinguish between clinical subtypes of stroke.

Though our analysis was aimed at understanding the genetic overlap between cardioembolicstroke andatrialfibrillation,we additionallyobserved genetic correlation between atrial fibrillation and undetermined stroke, a finding not observed in a previous investigation of atrial fibrillation PRS in ischemic stroke subtypes, albeit in a smaller sample.^28^ Perhaps contrary to expectation, we specifically found the atrial fibrillation polygenic risk score to be more strongly associated with the subset of etiology-undetermined strokes with an incomplete clinical evaluation, as opposed to those with cryptogenic stroke of a presumed, but not demonstrated, embolic source. These associations could be due to physician biases in diagnostic workups, rather than supporting a low prevalence of occult atrial fibrillation in presumed embolic strokes of undetermined source. Identifying stroke patients with atrial fibrillation is an important clinical challenge, as occult atrial fibrillation is well-known to cause strokes,^35,36^ and since such patients are at high risk for recurrent stroke, which is preventable with anticoagulation.^37,38^ Together, our findings indicate that atrial fibrillation genetic risk may augment clinical algorithms to determine stroke etiology, but will require further study.

The work presented here benefits from a number of improvements, including increased sample size; analysis of samples from a multicenter consortium, potentially enhancing the generalizability of the findings; and use of the CCS subtyping system, which provides more nuanced phenotyping, particularly in the cryptogenic subtype. Nevertheless, some limitations remain. Stroke is a heterogeneous condition that occurs later in life and has high lifetime prevalence (>15%^39^), features that can reduce statistical power. Further, sample sizes have lagged behind other GWAS efforts, a challenge further compounded by subtyping (nearly one-third of all cases are categorized as undetermined^23^). Reduced sample sizes impact power for discovery and make other analytic approaches – such as standard approaches for measuring trait correlation^16^ – unfeasible. Also, our sample is primarily comprised of Euroepan-ancestry samples, and work in non-Europeans, particularly in Africanancestry samples where risk of stroke is double that of European samples, is crucial. Finally, the current analysis does not analyze rare variation, which also likely contributes to disease susceptibility.^5^

We have shown that the cumulative genetic risk for atrial fibrillation in individuals with a stroke is similar to that reported in a larger population-based cohort.^25^ Genome-wide variation related to atrial fibrillation is substantially associated with cardioembolic stroke risk. Moreover, atrial fibrillation genetic risk was specific for cardioembolic stroke, and was not associated with the other primary stroke subtypes. The observation that atrial fibrillation genetic risk associated with strokes of undetermined cause supports the notion that undetected atrial fibrillation underlies a proportion of stroke risk in these individuals. Further work will need to incorporate emerging discoveries of rare genetic variants in atrial fibrillation, and explore the potential for genetic risk tools, including PRS’s performed via clinical-grade genotyping, to assist in the diagnostic workup of individuals with ischemic stroke.

## Acknowledgements

This work uses data from the National Institute of Neurological Disorders and Stroke - Stroke Genetics Network (NINDS-SiGN) Consortium and the Atrial Fibrillation Genetics (AFGen) Consortium. Members of NINDS-SiGN, the International Stroke Genetics Consortium, and AFGen are provided as an appendix to this paper.

Drs. Pulit, Mitchell, McArdle, and Kittner are supported by NIH grant R01NS100178. The NINDS-SiGN Consortium is supported by the NIH grants R01NS100178 and U01NS069208.

Dr. Pulit is supported by Veni Fellowship 016.186.071 (ZonMW) from the Dutch Organization for Scientific Research (Nederlandse Organisatie voor Wetenschappelijk Onderzoek, NWO).

Dr. Anderson is supported in part by K23NS086873, R01NS103924, an American Heart Association Strategically Focused Research Network in Atrial Fibrillation Award, and a Massachusetts General Hospital Center for Genomic Medicine Catalyst Award.

Dr. Lubitz is supported by NIH grants K23HL114724, an American Heart Association Strategically Focused Research Network in Atrial Fibrillation Award, and a Doris Duke Charitable Foundation Clinical Scientist Development Award 2014105.

Dr. Weng is supported by an American Heart Association Postdoctoral Fellowship Award 17POST33660226.

Drs. Ellinor and Benjamin are supported by the NIH grants R01HL092577 and R01HL128914, and an American Heart Association Strategically Focused Research Network in Atrial Fibrillation Award. Dr. Benjamin is additionally supported by the NIH grants 1RC1HL101056 and 1R01HL102214. Dr. Ellinor is additionally supported by the NIH grants R01HL104156 and K24HL105780; the National Heart, Lung, and Blood Institute (NHLBI); American Heart Association Established Investigator Award 13EIA14220013; and the Fondation Leducq 14CVD01.

## Shared Genetic Contributions to Atrial Fibrillation and Ischemic Stroke Risk

**Code and data release**

For access to information related to this project, including code, sample identifiers, SNP identifiers, links to summary-level data, and SNP weights used in the construction of the polygenic risk score, please see this GitHub repository: https://github.com/UMCUGenetics/Afib-Stroke-Overlap.

## Supplementary Figures

**Supplementary Figure 1.**
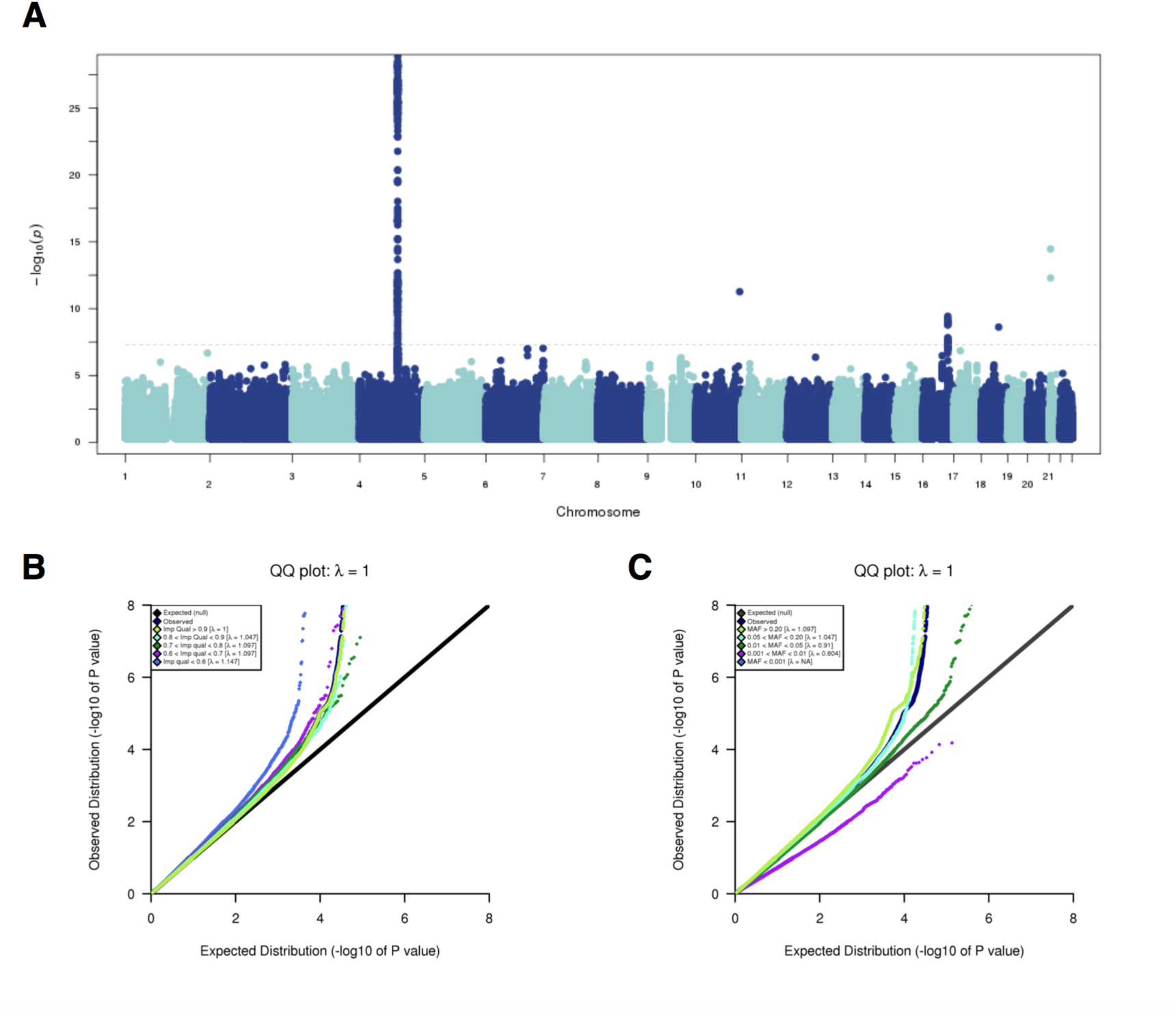
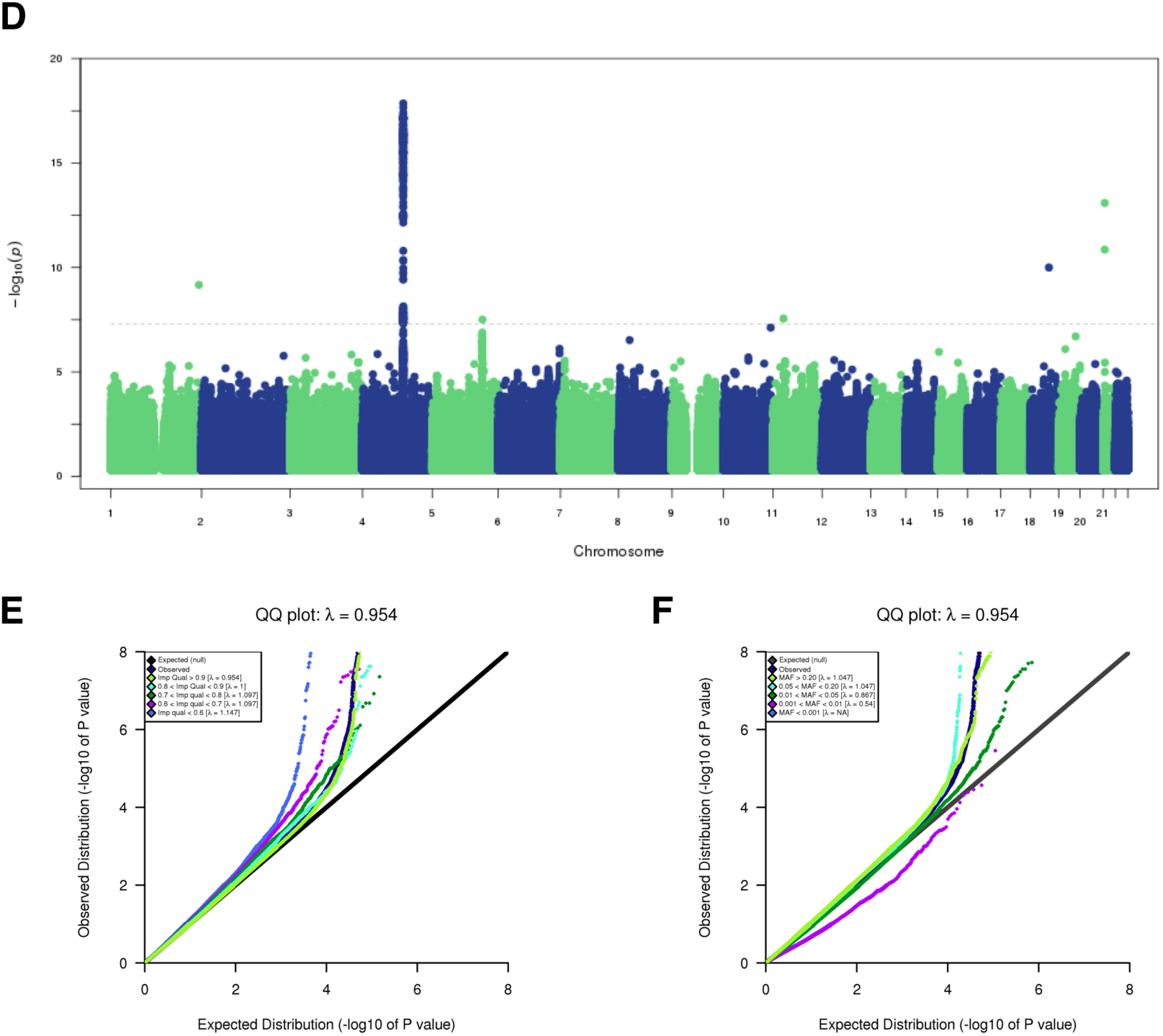
Genome-wide association study (GWAS) of atrial fibrillation in SiGN. (A) We performed a GWAS of 3,190 cases with atrial fibrillation, or paroxysmal atrial fibrillation, as well as other diagnoses suggestive of underlying atrial fibrillation, including left atrial thrombus, sick sinus syndrome, and atrial flutter. We additionally included 28,026 referents. We used a linear mixed model and adjusted the model for principal components and sex. The majority of atrial fibrillation risk loci identified through previous GWAS efforts were identified here at nominal significance or better (see **Supplementary Table 2**). The Manhattan plot only shows QC-passing SNPs with minor allele frequency > 1% and imputation quality score > 0.8. (B) Quantile-quantile (QQ) plot indicating SNPs stratified by minor allele frequency and the corresponding genomic inflation factor (lambda, λ) for each stratum. (C) QQ plot showing SNPs stratified by imputation quality and the corresponding lambda for each stratum. Figures D-F are identical to those of A-C, but for the analysis performed in atrial fibrillation cases only (N = 1,751). We performed this is an internal sensitivity analysis only, to ensure that more broadly defining the atrial fibrillation phenotype was not introducing additional phenotypic noise.

**Supplementary Figure 2.**
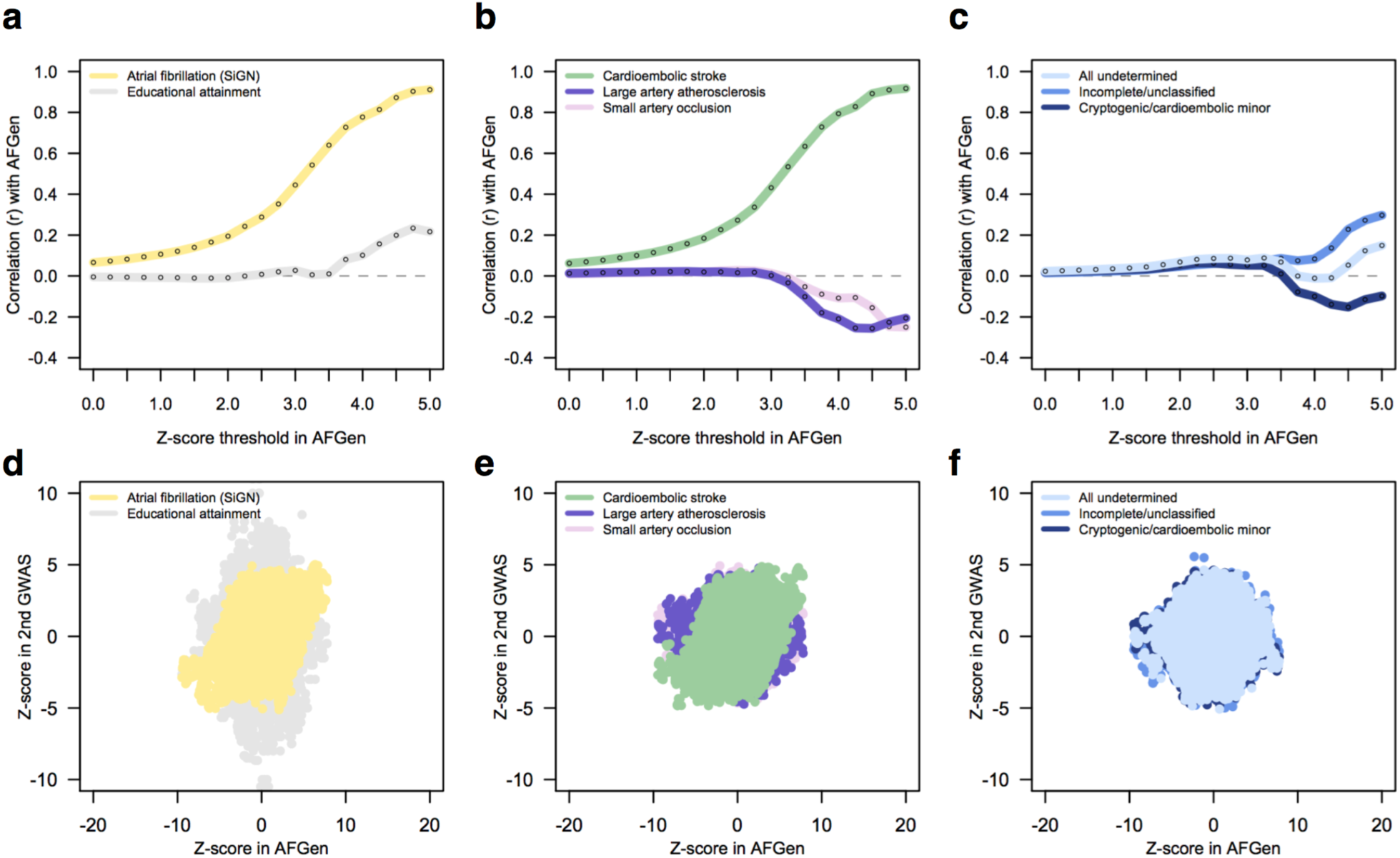
Genetic correlations between atrial fibrillation and ischemic stroke subtypes. To estimate genetic correlation between atrial fibrillation and ischemic stroke subtypes, we calculated Pearson’s r between SNP z-scores in the AFGen GWAS of atrial fibrillation and in GWAS of ischemic stroke subtypes and atrial fibrillation performed here in the SiGN data. Here, we present data identical to that shown in Figure 2 of the main manuscript, but removing ±2Mb around the two most significant loci discovered in atrial fibrillation and cardioembolic stroke: the region around *PITX2* (chromosome 4) and the region around *ZFHX3* (chromosome 16). (a) Genome wide, atrial fibrillation in AFGen and in SiGN correlate with increasing strength as the z-score in AFGen increases. Educational attainment is included here as a null comparator. (b) Genetic signal in cardioembolic stroke also correlates strongly with atrial fibrillation genetic signal in AFGen, but we do not observe correlation between atrial fibrillation and the other primary stroke subtypes. (c) Removing the *PITX2* and *ZFHX3* regions leaves only somewhat modest correlation between the incomplete/unclassified undetermined subtype and atrial fibrillation. Panels (d-f) show underlying data. Correlations restricted to those SNPs used in the polygenic risk score for atrial fibrillation were: AFGen vs atrial fibrillation in SiGN, r = 0.78; AFGen vs. cardioembolic stroke in SiGN, r = 0.75.

**Supplementary Figure 3.**
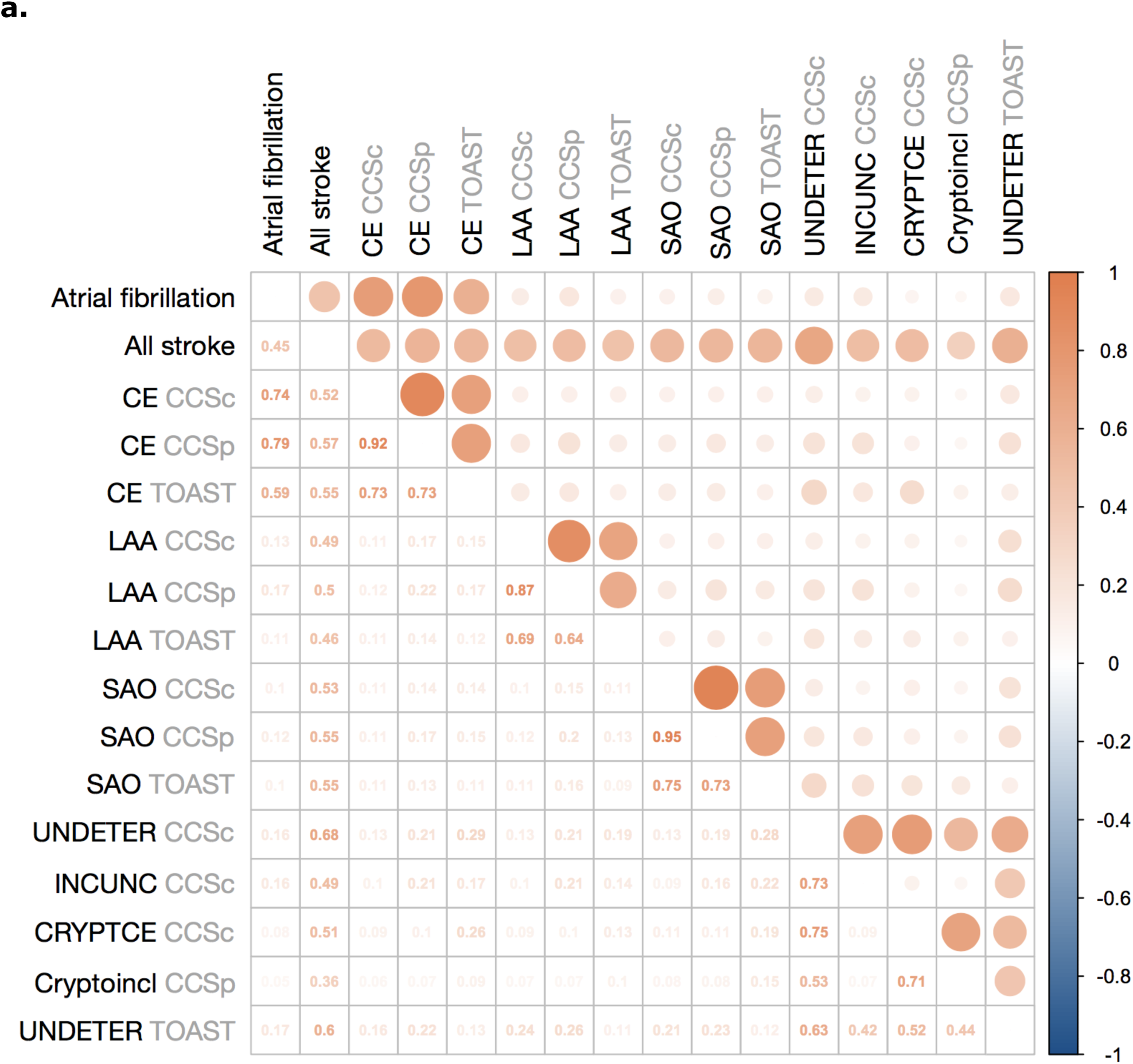
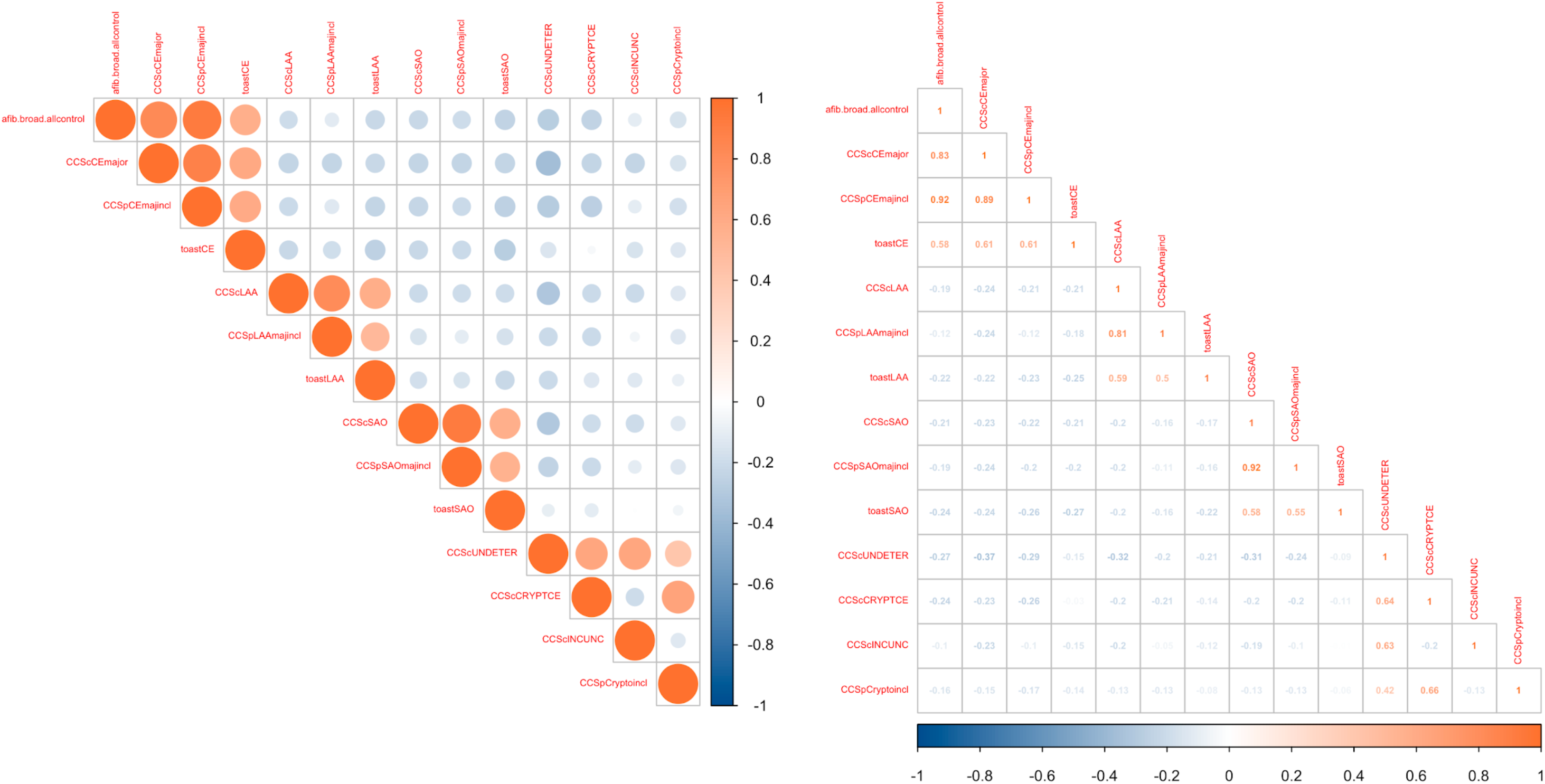
Genetic correlation and phenotypic correlation of atrial fibrillation and stroke subtypes in SiGN. (a) Using genome-wide SNP effects extracted from GWAS of atrial fibrillation, all stroke, and stroke subtypes, we calculated the Pearson’s correlation (r) between each pair of available phenotypes (blue indicates strong negative correlation; orange indicates strong positive correlation). Here, we show all correlations. Correlations are indicated by circle size in the upper half of the square, and the exact correlation values are shown in the lower half of the square. CE, cardioembolic stroke; LAA, large artery atherosclerosis; SAO, small artery occlusion; UNDETER, undetermined; INCUNC, incomplete/unclassified; CRYPTCE, cryptogenic and CE minor; Cryptoincl, cryptogenic; CCSc, CCS Causative subtyping system; CCSp, CCS Phenotypic subtyping system; TOAST, TOAST subtyping system. **b.** Same correlation calculations as in (a), but this time using the phenotypic data only (and looking in cases only, as all controls have the same phenotype). Note that the atrial fibrillation phenotypes and cardioembolic stroke phenotypes are highly correlated in the SiGN data (r = 0.83 between atrial fibrillation and cardioembolic stroke as determined by the CCS Causative subtype system). CE, cardioembolic stroke; LAA, large artery atherosclerosis; SAO, small artery occlusion; UNDETER, undetermined; INCUNC, incomplete/unclassified; CRYPTCE, cryptogenic and CE minor; Cryptoincl, cryptogenic; CCSc, CCS Causative subtyping system; CCSp, CCS Phenotypic subtyping system; TOAST, TOAST subtyping system.

**Supplementary Figure 4.**
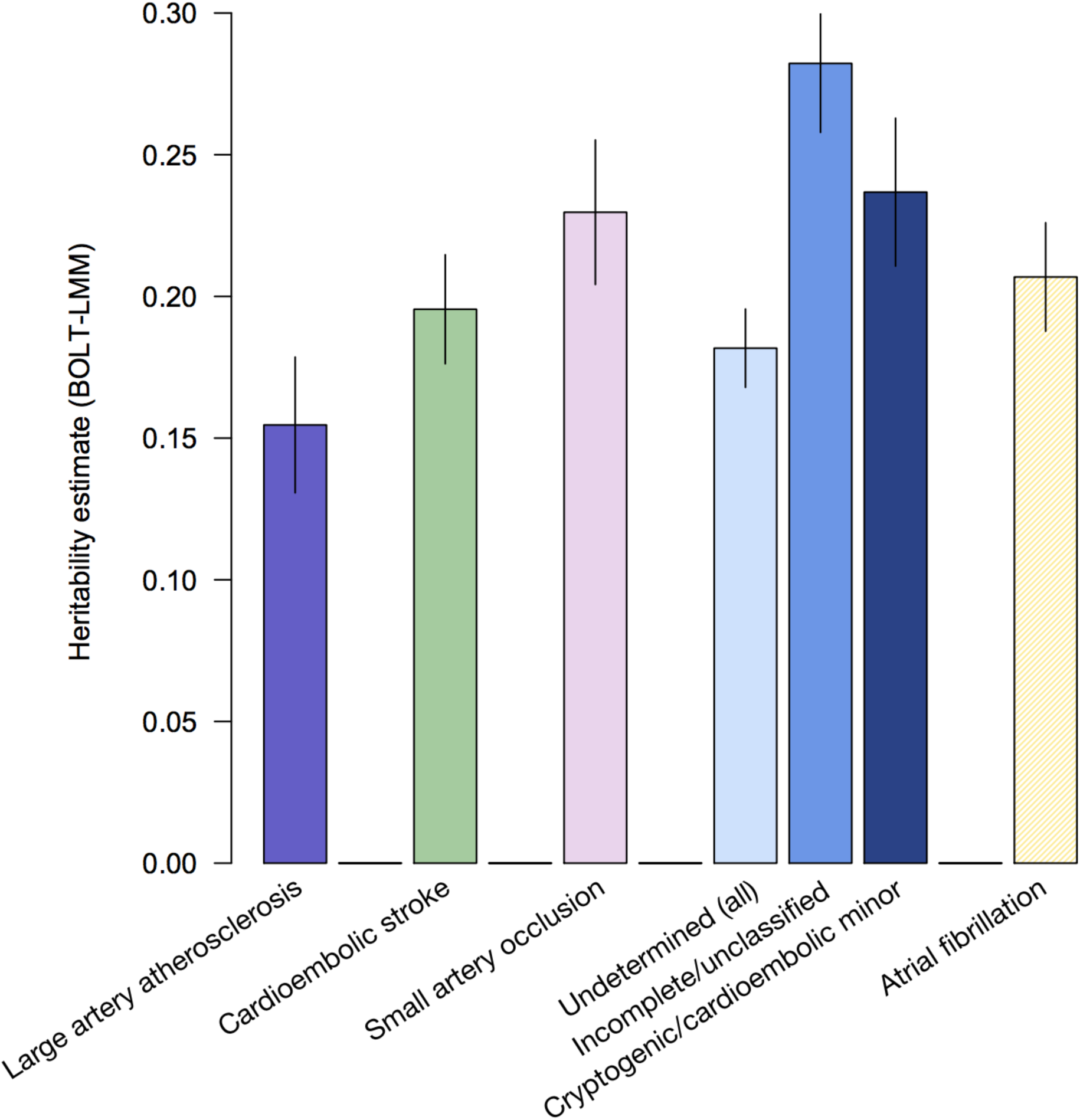
Estimated heritability of ischemic stroke subtypes and atrial fibrillation. Using all available stroke cases in SiGN, we estimated SNP-based heritability of the ischemic stroke subtypes (as sub-typed by the CCS Causative subtyping system) and atrial fibrillation (using the subset of 3,190 cases with atrial fibrillation) using BOLT-LMM and a genetic relationship matrix of high-quality SNPs converted to best-guess genotypes (imputation quality > 0.8, minor allele frequency > 0.01, and pruned at a linkage disequilibrium threshold of 0.2). We assumed a trait prevalence of 1% for all phenotypes. We found heritability estimates in cardioembolic stroke (green) and atrial fibrillation (yellow) to be approximately similar.

**Supplementary Figure 5.**
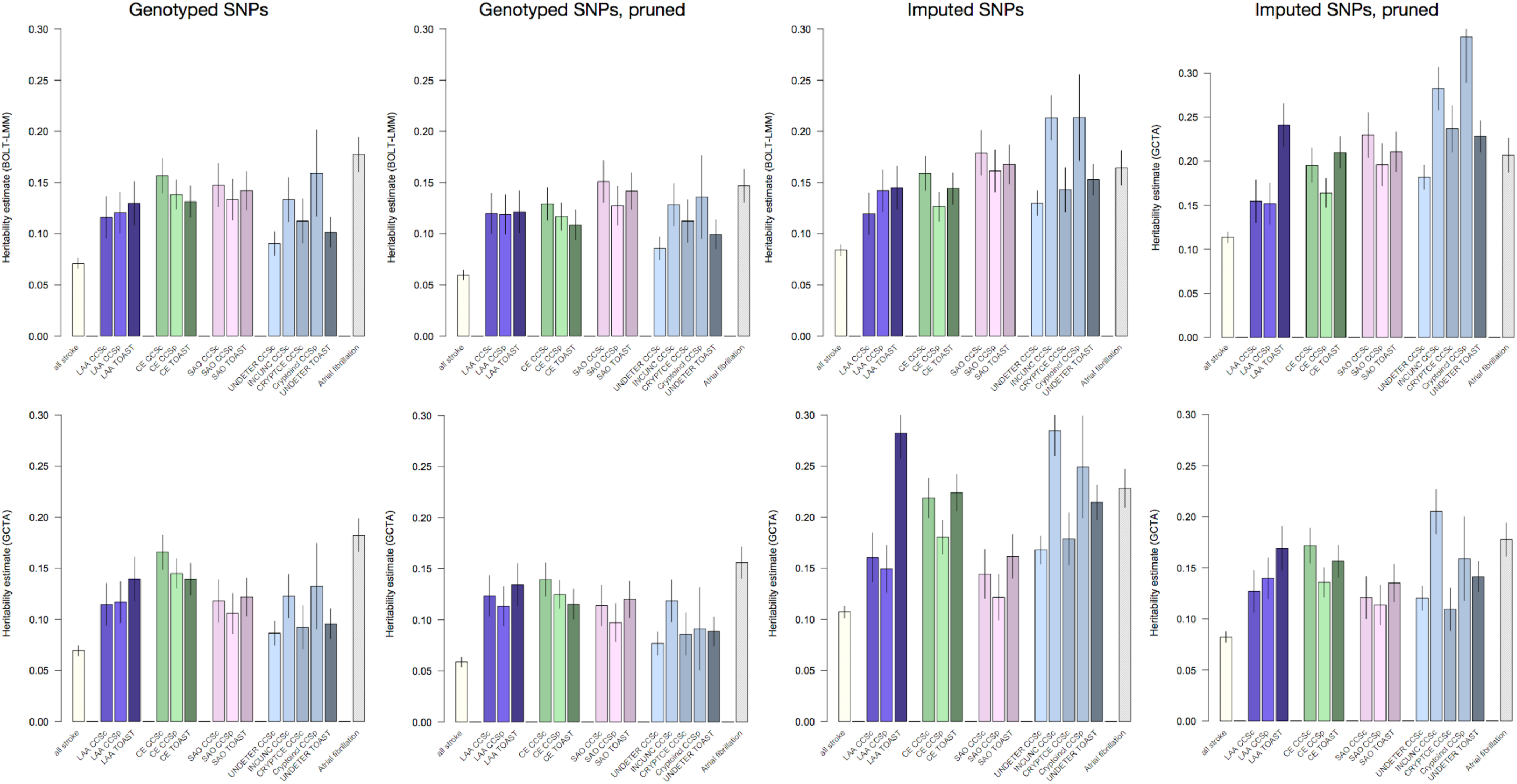
Heritability of ischemic stroke, its subtypes, and atrial fibrillation. We computed the SNP-based heritability of all stroke, all stroke subtypes, and atrial fibrillation using BOLT-LMM (top row) and GCTA (bottom row). All SNPs used for analysis had a minor allele frequency > 1% and imputation quality > 0.8 (for imputed SNPs). Imputed SNPs were converted to best-guess genotypes. We assumed a trait prevalence of 1% for all phenotypes and tested the robustness of 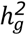 estimates to SNPs included in the GRM by using four different GRMs: (a) genotyped SNPs only; (b) genotyped, pruned, and filtered (see **Supplemental Methods**); (c) imputed; and (d) imputed, pruned, and filtered. We converted the imputed SNPs to hard-call genotypes before performing heritability analyses. Estimates are shown below, including error bars. The underlying data for these figures are provided in **Supplementary Table 3**. LAA, large artery atherosclerosis; CE, cardioembolic stroke; SAO, small artery occlusion; UNDETER, undetermined; INCUNC, incomplete/unclassified; CRYPTCE, cryptogenic and CE minor; Cryptoincl, cryptogenic; CCSc, CCS Causative subtyping system; CCSp, CCS Phenotypic subtyping system; TOAST, TOAST subtyping system.

**Supplementary Figure 6.**
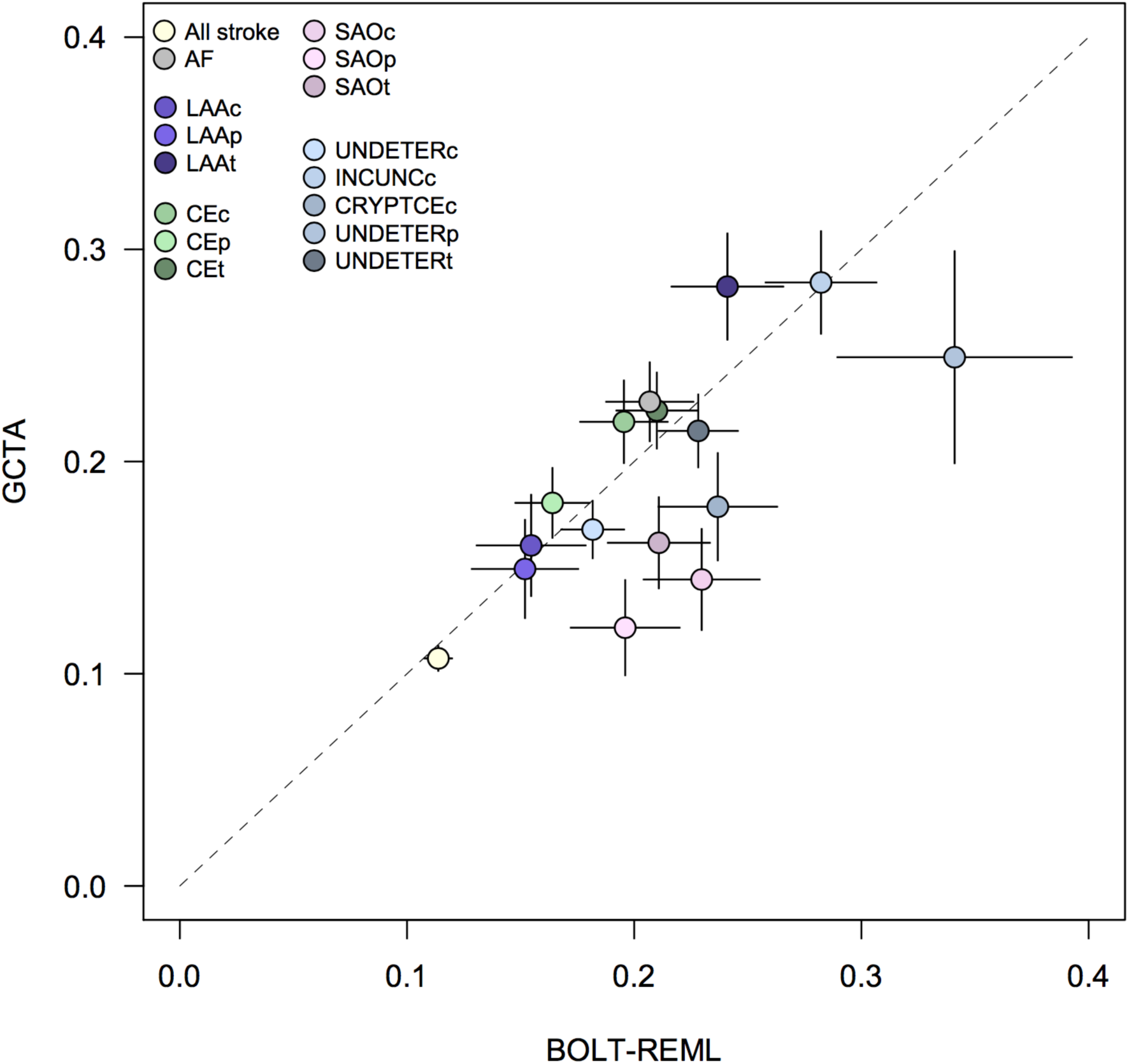
Comparison of heritability estimates from BOLT-LMM and GCTA. We computed the heritability of all stroke, all stroke subtypes, and atrial fibrillation using BOLT-LMM and GCTA, as shown in **Supplementary Figure 2**. Below, you will find a comparison of the two methods, with BOLT-REML on the x-axis and GCTA estimates on the y-axis. Error bars are shown for the respective estimates. AF, atrial fibrillation; CE, cardioembolic stroke; LAA, large artery atherosclerosis; SAO, small artery occlusion; UNDETER, undetermined; INCUNC, incomplete/unclassified; CRYPTCE, cryptogenic/CE minor; c, CCS Causative; p, CCS Phenotypic; t, TOAST.

**Supplementary Figure 7.**
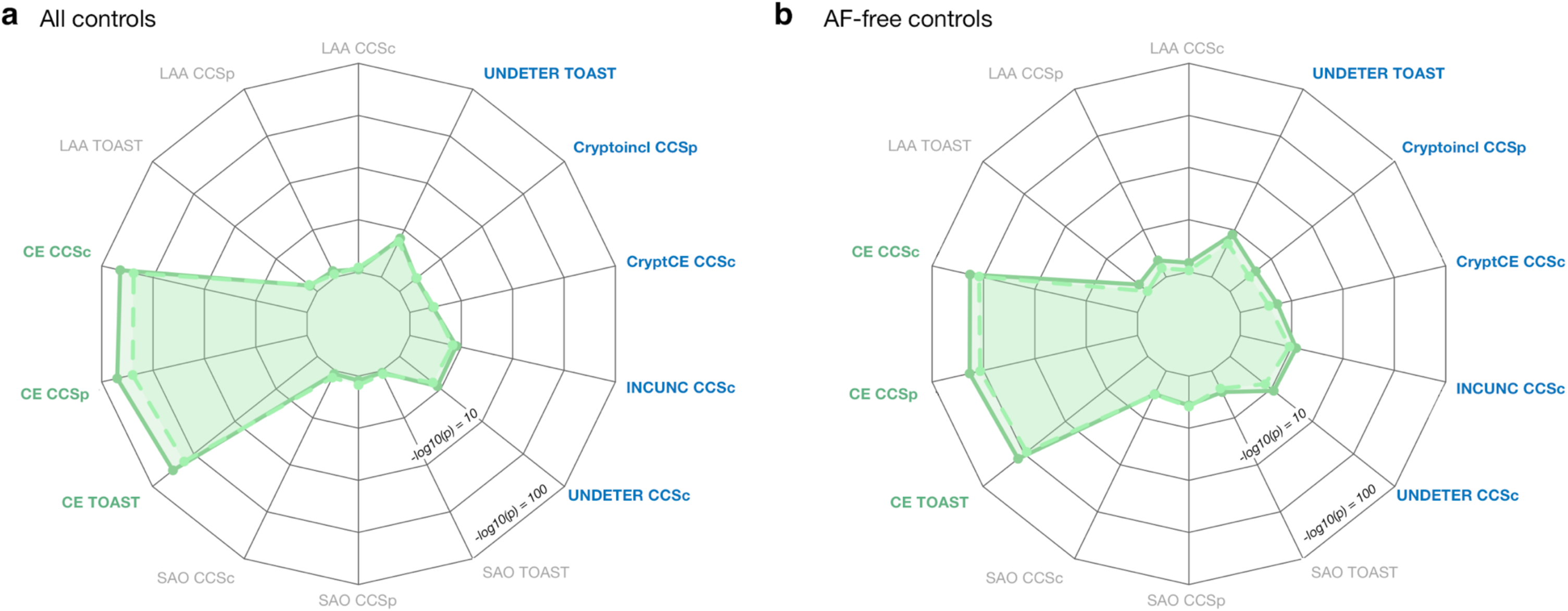
Association of atrial fibrillation polygenic risk score in ischemic stroke subtypes. We constructed a polygenic risk score (PRS) from atrial fibrillation-associated SNPs, and tested for association between the score and ischemic stroke subtypes using (a) all available controls (N = 28,026) and (b) controls without atrial fibrillation (N = 3,861). All subtypes from all available subtyping systems are shown here. The PRS strongly associated to cardioembolic stroke (subtypes highlighted in green font) in both sets of controls. In the atrial fibrillation-free set of controls (b) we observed nominal association of the PRS to incomplete/unclassified stroke. Undetermined subtypes are indicated in blue font. CE, cardioembolic stroke; LAA, large artery atherosclerosis; SAO, small artery occlusion; UNDETER, undetermined; INCUNC, incomplete/unclassified; CRYPTCE, cryptogenic and CE minor; Cryptoincl, cryptogenic; CCSc, CCS Causative subtyping system; CCSp, CCS Phenotypic subtyping system; TOAST, TOAST subtyping system.

## Supplementary Tables

**Supplementary Table 1.**
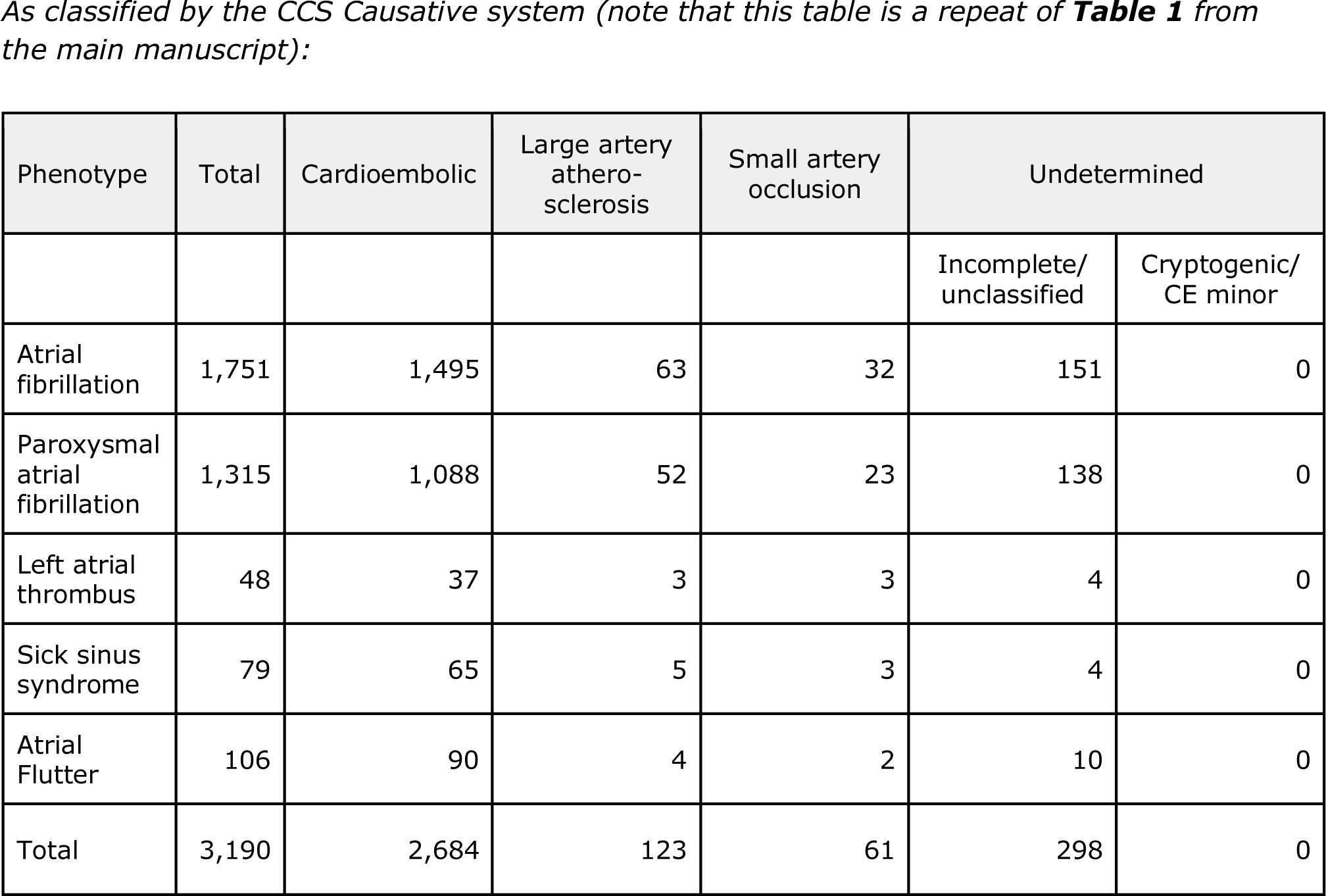
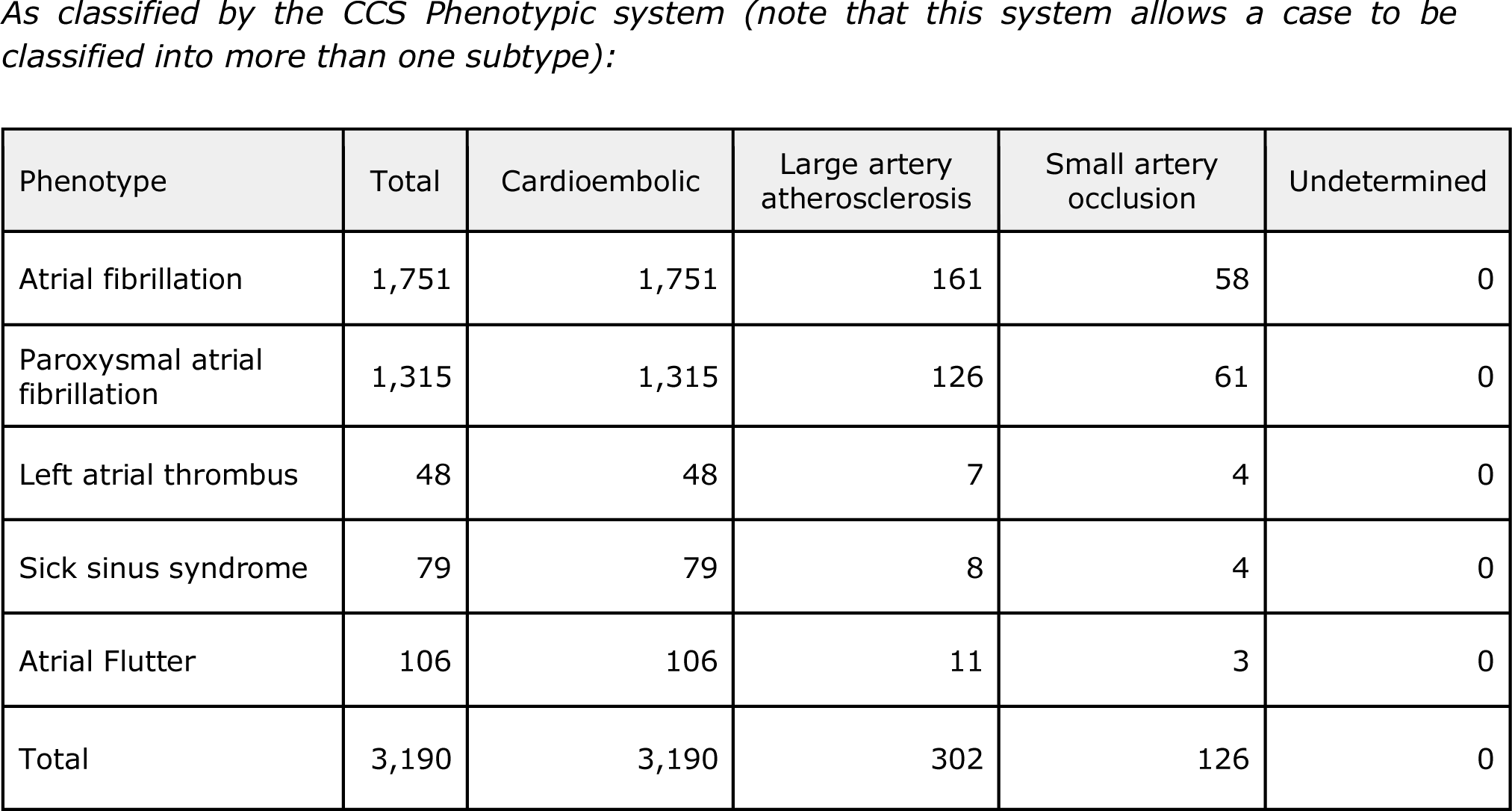
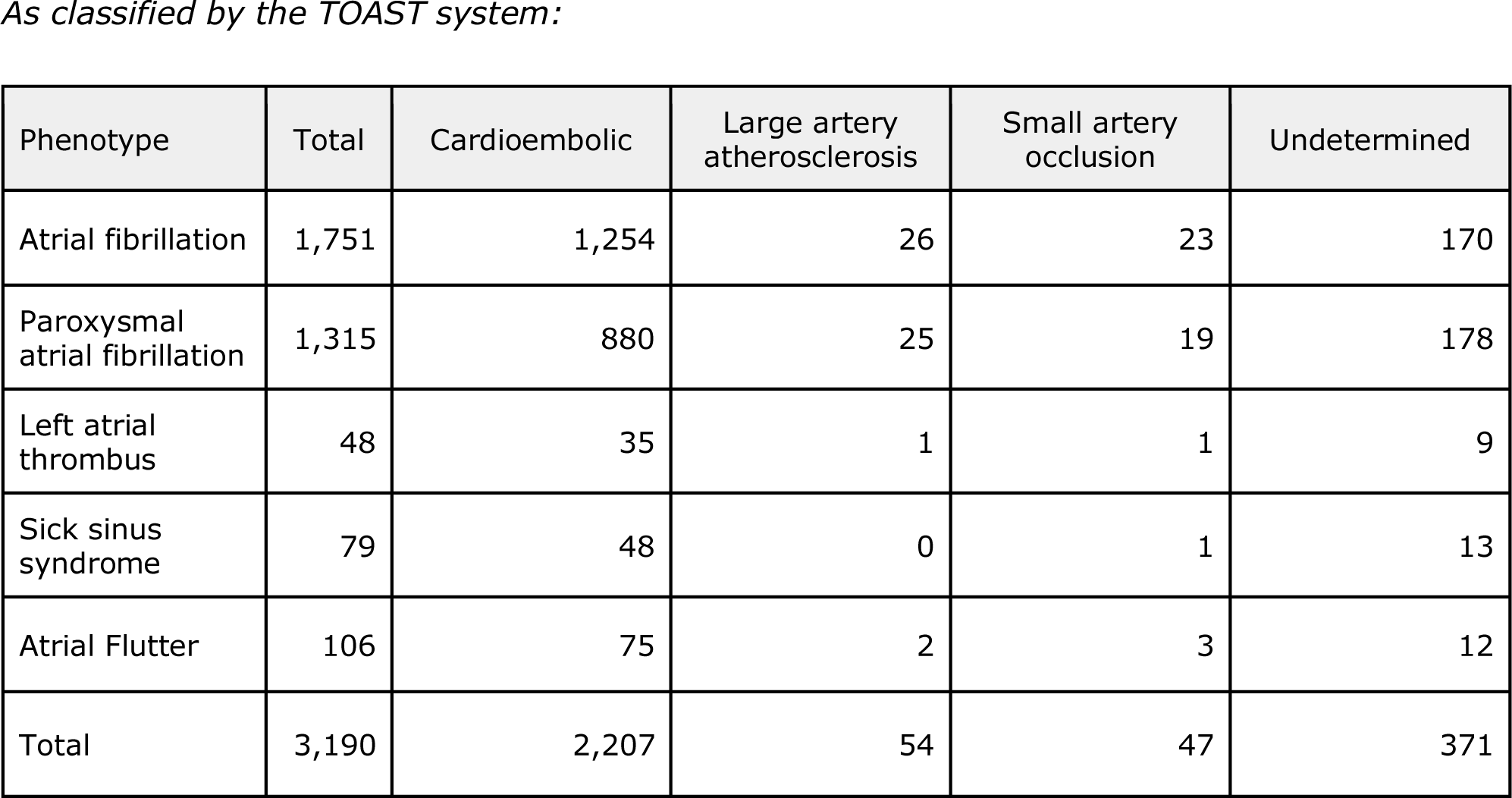
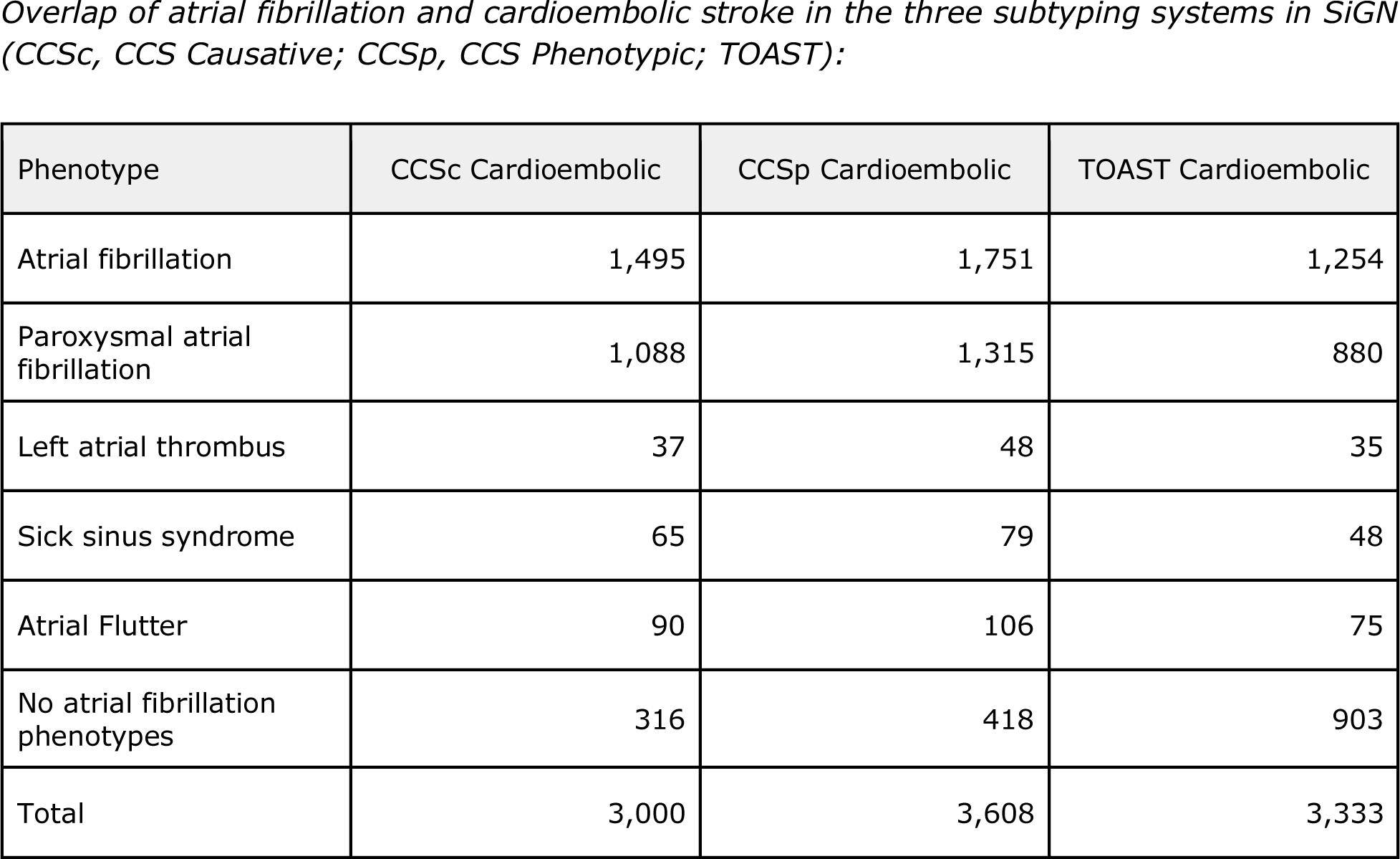
Atrial fibrillation cases and controls available from the Stroke Genetics Network (SiGN) Consortium.

| Phenotype | Total | Cardioembolic | Large artery atherosclerosis | Small artery occlusion | Undetermined |  |
| --- | --- | --- | --- | --- | --- | --- |
|  |  |  |  |  | Incomplete/unclassified | Cryptogenic/CE minor |
| Atrial fibrillation | 1,751 | 1,495 | 63 | 32 | 151 | 0 |
| Paroxysmal atrial fibrillation | 1,315 | 1,088 | 52 | 23 | 138 | 0 |
| Left atrial thrombus | 48 | 37 | 3 | 3 | 4 | 0 |
| Sick sinus syndrome | 79 | 65 | 5 | 3 | 4 | 0 |
| Atrial Flutter | 106 | 90 | 4 | 2 | 10 | 0 |
| Total | 3,190 | 2,684 | 123 | 61 | 298 | 0 |

| Phenotype | Total | Cardioembolic | Large artery atherosclerosis | Small artery occlusion | Undetermined |
| --- | --- | --- | --- | --- | --- |
| Atrial fibrillation | 1,751 | 1,751 | 161 | 58 | 0 |
| Paroxysmal atrial fibrillation | 1,315 | 1,315 | 126 | 61 | 0 |
| Left atrial thrombus | 48 | 48 | 7 | 4 | 0 |
| Sick sinus syndrome | 79 | 79 | 8 | 4 | 0 |
| Atrial Flutter | 106 | 106 | 11 | 3 | 0 |
| Total | 3,190 | 3,190 | 302 | 126 | 0 |

*As classified by the TOAST system:*
| Phenotype | Total | Cardioembolic | Large artery atherosclerosis | Small artery occlusion | Undetermined |
| --- | --- | --- | --- | --- | --- |
| Atrial fibrillation | 1,751 | 1,254 | 26 | 23 | 170 |
| Paroxysmal atrial fibrillation | 1,315 | 880 | 25 | 19 | 178 |
| Left atrial thrombus | 48 | 35 | 1 | 1 | 9 |
| Sick sinus syndrome | 79 | 48 | 0 | 1 | 13 |
| Atrial Flutter | 106 | 75 | 2 | 3 | 12 |
| Total | 3,190 | 2,207 | 54 | 47 | 371 |

| Phenotype | CCSc Cardioembolic | CCSp Cardioembolic | TOAST Cardioembolic |
| --- | --- | --- | --- |
| Atrial fibrillation | 1,495 | 1,751 | 1,254 |
| Paroxysmal atrial fibrillation | 1,088 | 1,315 | 880 |
| Left atrial thrombus | 37 | 48 | 35 |
| Sick sinus syndrome | 65 | 79 | 48 |
| Atrial Flutter | 90 | 106 | 75 |
| No atrial fibrillation phenotypes | 316 | 418 | 903 |
| Total | 3,000 | 3,608 | 3,333 |

**Supplementary Table 2.**
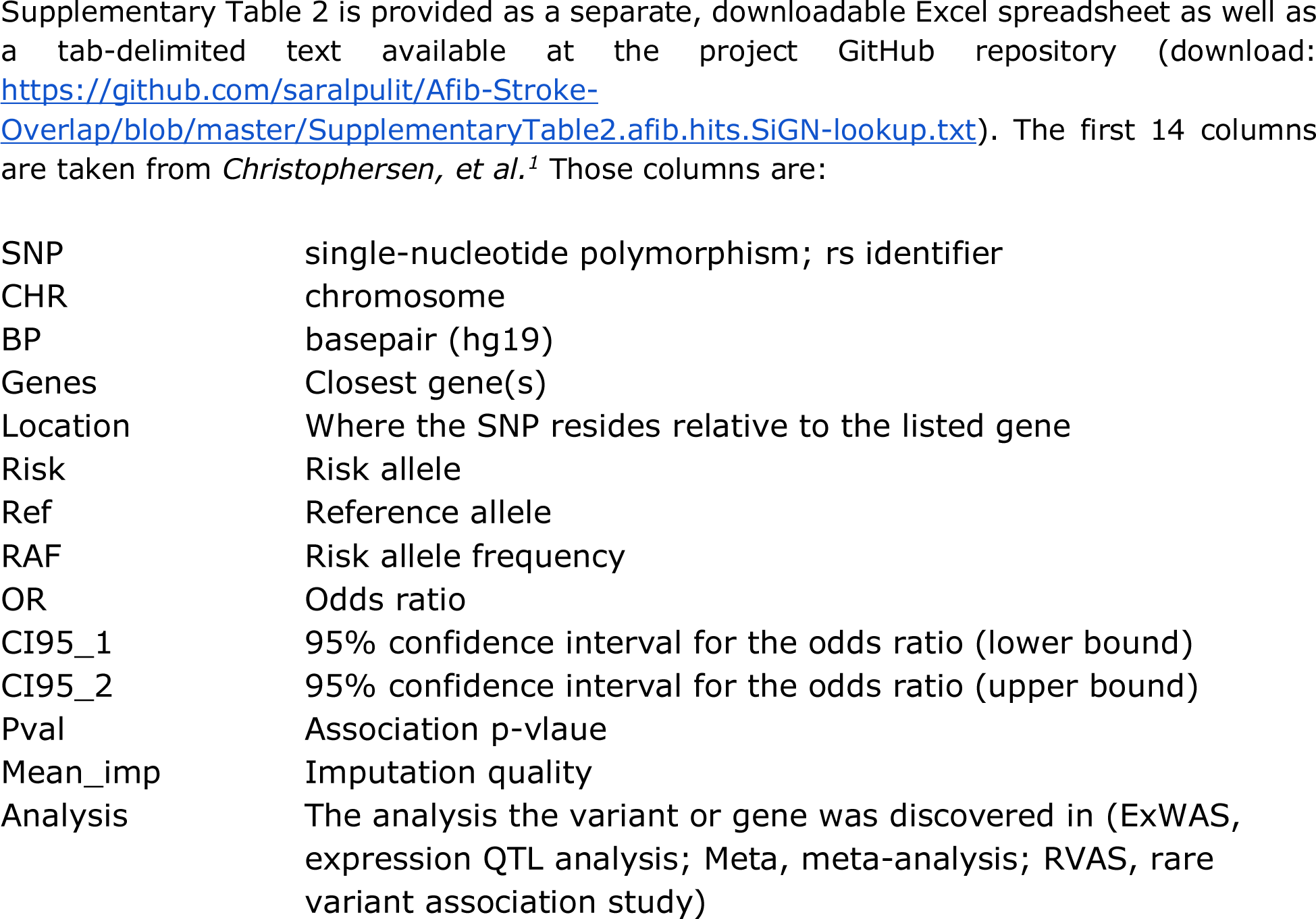
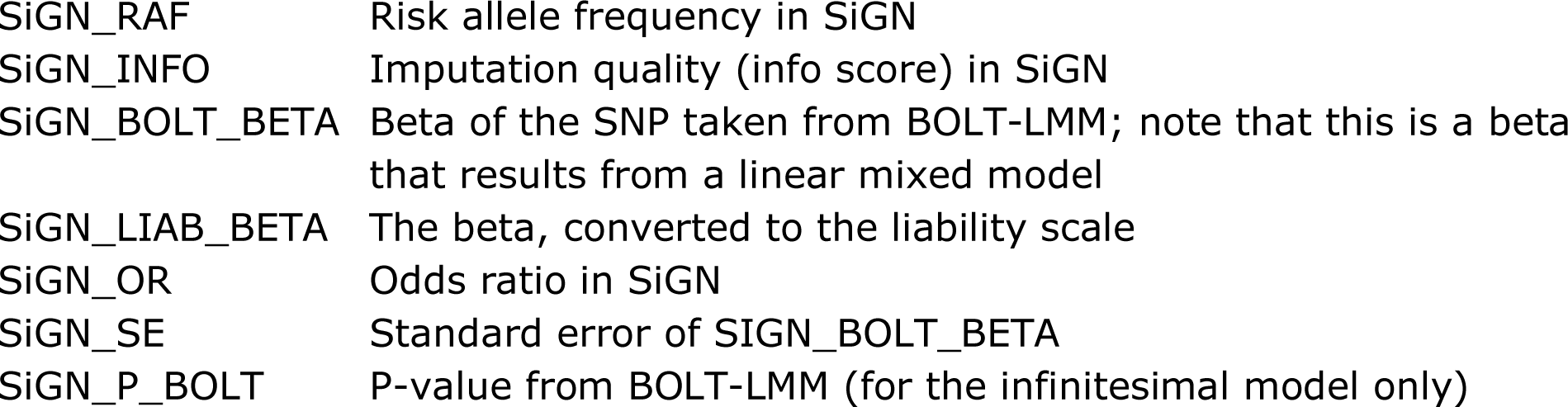
Look-up of previously-associated atrial fibrillation SNPs in SiGN. After performing a GWAS of atrial fibrillation in the SiGN data, we looked up the 26 known genetic risk loci for atrial fibrillation, as identified in the latest GWAS.^1^ Twenty-four of the 25 signals present in the SiGN data were directionally consistent with the previous GWAS. The only signal not directionally consistent was discovered through eQTL analysis. One signal, a rare variant burden signal, was absent from our data (all SNPs here have allele frequency > 1%). The remaining columns provided are data points extracted from the atrial fibrillation GWAS in SiGN. They are:

|  |  |
| --- | --- |
| SNP | single-nucleotide polymorphism; rs identifier |
| CHR | chromosome |
| BP | basepair (hg19) |
| Genes | Closest gene(s) |
| Location | Where the SNP resides relative to the listed gene |
| Risk | Risk allele |
| Ref | Reference allele |
| RAF | Risk allele frequency |
| OR | Odds ratio |
| CI95_1 | 95% confidence interval for the odds ratio (lower bound) |
| CI95_2 | 95% confidence interval for the odds ratio (upper bound) |
| Pval | Association p-value |
| Mean_imp | Imputation quality |
| Analysis | The analysis the variant or gene was discovered in (ExWAS, expression QTL analysis; Meta, meta-analysis; RVAS, rare variant association study) |
| SiGN_RAF | Risk allele frequency in SiGN |
| SiGN_INFO | Imputation quality (info score) in SiGN |
| SiGN_BOLT_BETA | Beta of the SNP taken from BOLT-LMM; note that this is a beta that results from a linear mixed model |
| SiGN_LIAB_BETA | The beta, converted to the liability scale |
| SiGN_OR | Odds ratio in SiGN |
| SiGN_SE | Standard error of SiGN_BOLT_BETA |
| SiGN_P_BOLT | P-value from BOLT-LMM (for the infinitesimal model only) |

**Supplementary Table 3.** Genetic correlations between atrial fibrillation and ischemic stroke subtypes. To estimate genetic correlation between atrial fibrillation and ischemic stroke subtypes, we calculated Pearson’s r between SNP z-scores in the Atrial Fibrillation Genetics (AFGen) GWAS of atrial fibrillation and in GWAS of ischemic stroke subtypes and atrial fibrillation performed here in the SiGN data. The correlation calculations are provided in this table, which is split into two parts and is available to download in text format here:

| Column | Definition |
| --- | --- |
| Z.threshold | Z-score threshold used to subset AFGen SNPs |
| EduYrs.Z | Correlation with z-scores from educational attainment GWAS |
| afib.broad.Z | Correlation with z-scores from atrial fibrillation (broadly defined phenotype) GWAS |
| allstroke.Z | Correlation with z-scores from all stroke GWAS |
| CCScCEmajor.Z | Correlation with z-scores from CCSc CE GWAS |
| CCScCRYPTCE.Z | Correlation with z-scores from CCSc CRYPTCE GWAS |
| CCScINCUNC.Z | Correlation with z-scores from CCSc INCUNC GWAS |
| CCScLAA.Z | Correlation with z-scores from CCSc LAA GWAS |
| CCScSAO.Z | Correlation with z-scores from CCSc SAO GWAS |
| CCScUNDETER.Z | Correlation with z-scores from CCSc UNDETER GWAS |
| CCSpCEmajincl.Z | Correlation with z-scores from CCSp CE GWAS |
| CCSpCryptoincl.Z | Correlation with z-scores from CCSp Cryptogenic GWAS |
| CCSpLAAmajincl.Z | Correlation with z-scores from CCSp LAA GWAS |
| CCSpSAOmajincl.Z | Correlation with z-scores from CCSp SAO GWAS |
| toastCE.Z | Correlation with z-scores from TOAST CE GWAS |
| toastLAA.Z | Correlation with z-scores from TOAST LAA GWAS |
| toastSAO.Z | Correlation with z-scores from TOAST SAO GWAS |
| toastUNDETER.Z | Correlation with z-scores from TOAST UNDETER GWAS |

CCSc, CCS Causative subtyping system; CCSp, CCS Phenotypic subtyping system; TOAST, TOAST subtyping system; CE, cardioembolic stroke; LAA, large artery atherosclerosis; SAO, small artery occlusion; UNDETER, undetermined; INCUNC, incomplete/unclassified; CRYPTCE, cryptogenic and CE minor.

**Supplementary Table 4.**
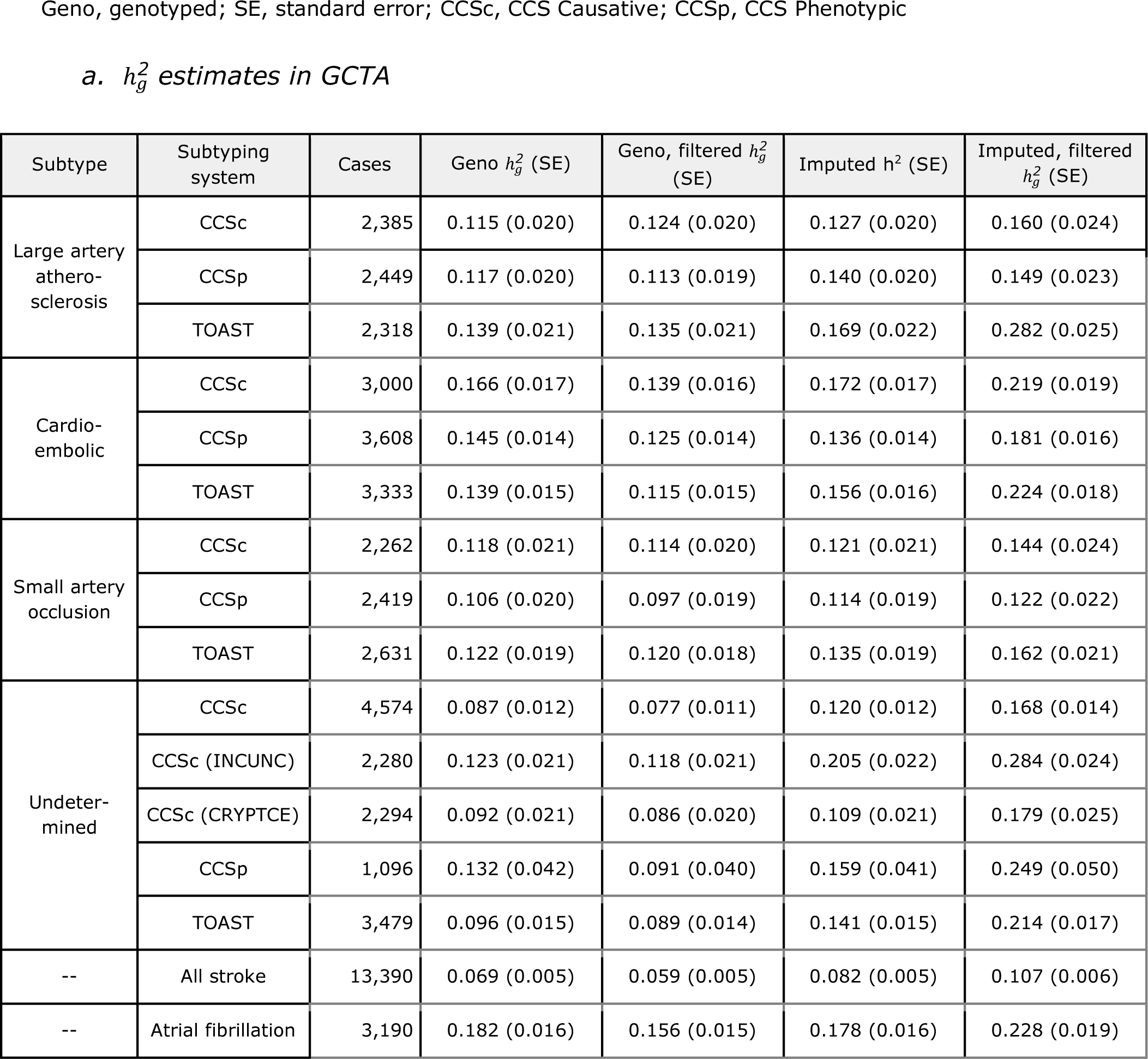
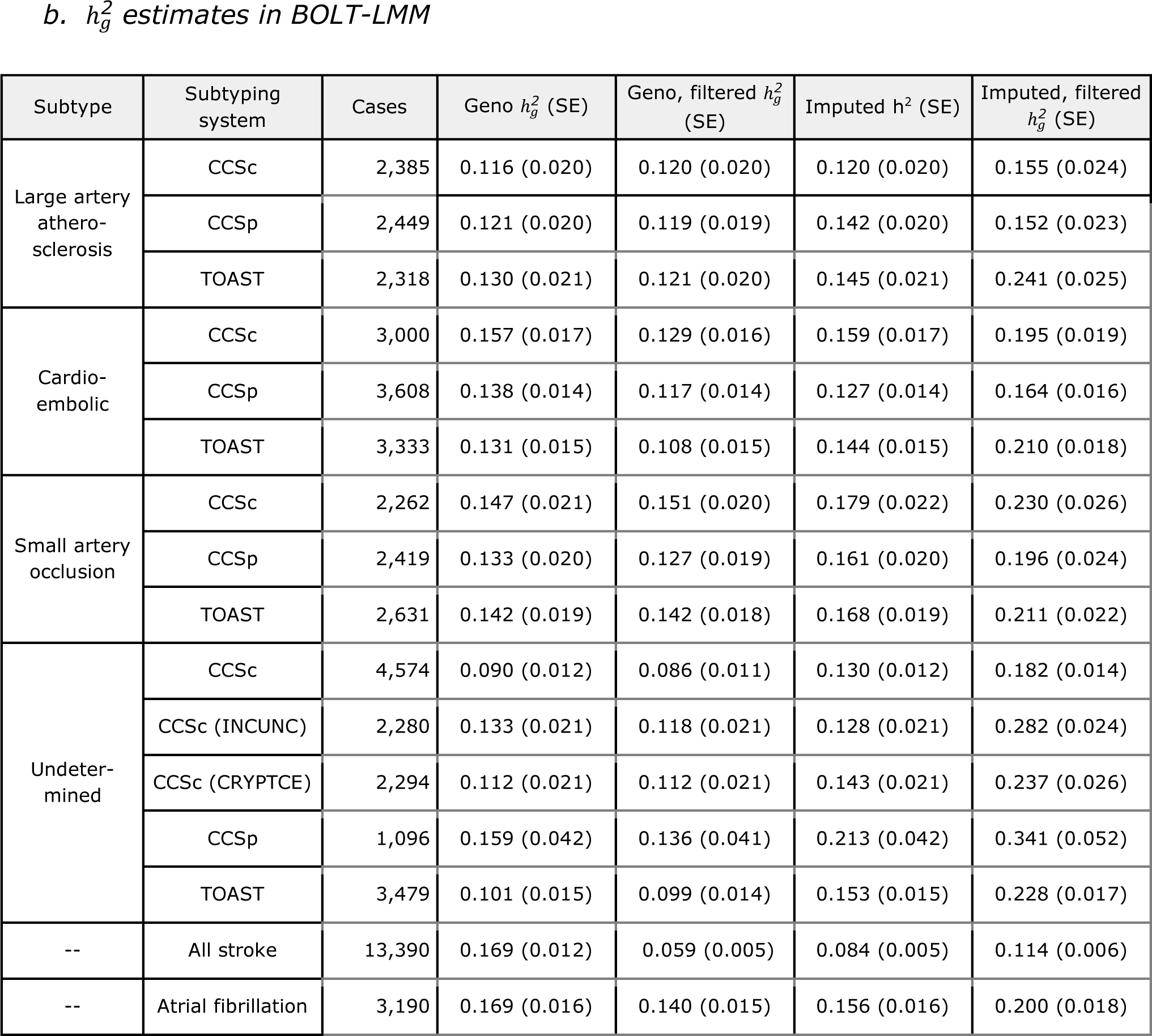
Heritability calculations in atrial fibrillation and ischemic stroke subtypes. (a) We calculated the SNP-based heritability 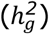 of atrial fibrillation, all ischemic stroke, and the stroke subtypes using GCTA^2^. All SNPs used had minor allele frequency > 1% and imputation quality > 0.8 (for imputed SNPs). Imputed SNPs were converted to best-guess genotypes. We assumed a trait prevalence of 1% for all phenotypes and tested the robustness of 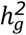.estimates to SNPs included in the GRM by using four different GRMs: (i) genotyped only; (ii) genotyped, pruned, and filtered (see **Supplemental Methods**); (iii) imputed; and (iv) imputed, pruned, and filtered. (b) We performed the exact same analysis but using BOLT-LMM to estimate.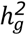 BOLT-LMM estimates were converted to the liability scale (see **Supplemental Methods**).

| Subtype | Subtyping system | Cases | Geno $h_g^2$ (SE) | Geno, filtered $h_g^2$ (SE) | Imputed $h^2$ (SE) | Imputed, filtered $h_g^2$ (SE) |
| --- | --- | --- | --- | --- | --- | --- |
| Large artery atherosclerosis | CCSc | 2,385 | 0.115 (0.020) | 0.124 (0.020) | 0.127 (0.020) | 0.160 (0.024) |
|  | CCSp | 2,449 | 0.117 (0.020) | 0.113 (0.019) | 0.140 (0.020) | 0.149 (0.023) |
|  | TOAST | 2,318 | 0.139 (0.021) | 0.135 (0.021) | 0.169 (0.022) | 0.282 (0.025) |
| Cardio-embolic | CCSc | 3,000 | 0.166 (0.017) | 0.139 (0.016) | 0.172 (0.017) | 0.219 (0.019) |
|  | CCSp | 3,608 | 0.145 (0.014) | 0.125 (0.014) | 0.136 (0.014) | 0.181 (0.016) |
|  | TOAST | 3,333 | 0.139 (0.015) | 0.115 (0.015) | 0.156 (0.016) | 0.224 (0.018) |
| Small artery occlusion | CCSc | 2,262 | 0.118 (0.021) | 0.114 (0.020) | 0.121 (0.021) | 0.144 (0.024) |
|  | CCSp | 2,419 | 0.106 (0.020) | 0.097 (0.019) | 0.114 (0.019) | 0.122 (0.022) |
|  | TOAST | 2,631 | 0.122 (0.019) | 0.120 (0.018) | 0.135 (0.019) | 0.162 (0.021) |
| Undetermined | CCSc | 4,574 | 0.087 (0.012) | 0.077 (0.011) | 0.120 (0.012) | 0.168 (0.014) |
|  | CCSc (INCUNC) | 2,280 | 0.123 (0.021) | 0.118 (0.021) | 0.205 (0.022) | 0.284 (0.024) |
|  | CCSc (CRYPTCE) | 2,294 | 0.092 (0.021) | 0.086 (0.020) | 0.109 (0.021) | 0.179 (0.025) |
|  | CCSp | 1,096 | 0.132 (0.042) | 0.091 (0.040) | 0.159 (0.041) | 0.249 (0.050) |
|  | TOAST | 3,479 | 0.096 (0.015) | 0.089 (0.014) | 0.141 (0.015) | 0.214 (0.017) |
| -- | All stroke | 13,390 | 0.069 (0.005) | 0.059 (0.005) | 0.082 (0.005) | 0.107 (0.006) |
| -- | Atrial fibrillation | 3,190 | 0.182 (0.016) | 0.156 (0.015) | 0.178 (0.016) | 0.228 (0.019) |
| Large artery atherosclerosis | CCSc | 2,385 | 0.116 (0.020) | 0.120 (0.020) | 0.120 (0.020) | 0.155 (0.024) |
|  | CCSp | 2,449 | 0.121 (0.020) | 0.119 (0.019) | 0.142 (0.020) | 0.152 (0.023) |
|  | TOAST | 2,318 | 0.130 (0.021) | 0.121 (0.020) | 0.145 (0.021) | 0.241 (0.025) |
| Cardio-embolic | CCSc | 3,000 | 0.157 (0.017) | 0.129 (0.016) | 0.159 (0.017) | 0.195 (0.019) |
|  | CCSp | 3,608 | 0.138 (0.014) | 0.117 (0.014) | 0.127 (0.014) | 0.164 (0.016) |
|  | TOAST | 3,333 | 0.131 (0.015) | 0.108 (0.015) | 0.144 (0.015) | 0.210 (0.018) |
| Small artery occlusion | CCSc | 2,262 | 0.147 (0.021) | 0.151 (0.020) | 0.179 (0.022) | 0.230 (0.026) |
|  | CCSp | 2,419 | 0.133 (0.020) | 0.127 (0.019) | 0.161 (0.020) | 0.196 (0.024) |
|  | TOAST | 2,631 | 0.142 (0.019) | 0.142 (0.018) | 0.168 (0.019) | 0.211 (0.022) |
| Undetermined | CCSc | 4,574 | 0.090 (0.012) | 0.086 (0.011) | 0.130 (0.012) | 0.182 (0.014) |
|  | CCSc (INCUNC) | 2,280 | 0.133 (0.021) | 0.118 (0.021) | 0.128 (0.021) | 0.282 (0.024) |
|  | CCSc (CRYPTCE) | 2,294 | 0.112 (0.021) | 0.112 (0.021) | 0.143 (0.021) | 0.237 (0.026) |
|  | CCSp | 1,096 | 0.159 (0.042) | 0.136 (0.041) | 0.213 (0.042) | 0.341 (0.052) |
|  | TOAST | 3,479 | 0.101 (0.015) | 0.099 (0.014) | 0.153 (0.015) | 0.228 (0.017) |
| -- | All stroke | 13,390 | 0.169 (0.012) | 0.059 (0.005) | 0.084 (0.005) | 0.114 (0.006) |
| -- | Atrial fibrillation | 3,190 | 0.169 (0.016) | 0.140 (0.015) | 0.156 (0.016) | 0.200 (0.018) |

**Supplementary Table 5.** Association between the atrial fibrillation polygenic risk score and ischemic stroke subtypes. We tested the association between a polygenic risk score (PRS) constructed from atrial fibrillation-associated SNPs and all stroke subtypes. The results of those association tests are shown here. We used two groups of controls: all available controls (N = 28,026 in the model without clinical covariates; N = 14,357 in the model with clinical covariates) and all controls that were free of atrial fibrillation (AF, N = 3,860 in the model without clinical covariates; N = 3,786 in the model with clinical covariates). All analyses were adjusted for sex and principal components (PCs). Regression analyses were optionally adjusted for clinical covariates (age, cardiovascular disease, type 2 diabetes status, smoking status, and hypertension).

| Case definition | Control definition | Cases |  | Logistic regression, adjusted for PCs and sex |  |  | Logistic regression, adjusted for PCs, sex, and clinical covariates |  |  |
| --- | --- | --- | --- | --- | --- | --- | --- | --- | --- |
|  |  | w/out clinical covars | with clinical covars | Beta | SE | P-value | Beta | SE | P-value |
| CCSc LAA | Non-AF controls | 2,385 | 2,093 | 0.008 | 0.015 | 0.600 | 0.002 | 0.018 | 0.929 |
| CCSc LAA | All controls | 2,385 | 2,093 | -0.002 | 0.012 | 0.885 | -0.004 | 0.013 | 0.786 |
| CCSp LAA | Non-AF controls | 2,449 | 2,149 | 0.016 | 0.016 | 0.315 | 0.010 | 0.018 | 0.570 |
| CCSp LAA | All controls | 2,449 | 2,149 | 0.004 | 0.011 | 0.694 | 0.002 | 0.013 | 0.850 |
| TOAST LAA | Non-AF controls | 2,318 | 1,884 | 0.010 | 0.016 | 0.528 | 0.000 | 0.018 | 0.980 |
| TOAST LAA | All controls | 2,318 | 1,884 | -0.006 | 0.012 | 0.594 | -0.008 | 0.014 | 0.550 |
| *Results after standardizing PRS to a z-score* |  |  |  |  |  |  |  |  |  |
| CCSc LAA | Non-AF controls | 2,385 | 2,093 | 0.016 | 0.030 | 0.600 | 0.003 | 0.035 | 0.929 |
| CCSc LAA | All controls | 2,385 | 2,093 | -0.003 | 0.022 | 0.885 | -0.007 | 0.026 | 0.786 |
| CCSp LAA | Non-AF controls | 2,449 | 2,149 | 0.031 | 0.030 | 0.315 | 0.020 | 0.035 | 0.570 |
| CCSp LAA | All controls | 2,449 | 2,149 | 0.009 | 0.022 | 0.694 | 0.005 | 0.026 | 0.850 |
| TOAST LAA | Non-AF controls | 2,318 | 1,884 | 0.019 | 0.031 | 0.528 | -0.001 | 0.036 | 0.980 |
| TOAST LAA | All controls | 2,318 | 1,884 | -0.012 | 0.023 | 0.594 | -0.016 | 0.027 | 0.550 |

| Case definition | Control definition (N) | Cases |  | Logistic regression, adjusted for PCs and sex |  |  | Logistic regression, adjusted for PCs, sex, and clinical covariates |  |  |
| --- | --- | --- | --- | --- | --- | --- | --- | --- | --- |
|  |  |  |  | Beta | SE | P-value | Beta | SE | P-value |
| CCSc CE | Non-AF (3,869) | 3,000 | 2,725 | 0.187 | 0.014 | **1.59E-42** | 0.218 | 0.018 | **1.40E-34** |
| CCSc CE | All (28,026) | 3,000 | 2,725 | 0.169 | 0.010 | **1.01E-65** | 0.173 | 0.012 | **1.45E-48** |
| CCSp CE | Non-AF (3,869) | 3,608 | 3,281 | 0.178 | 0.013 | **6.98E-43** | 0.203 | 0.017 | **8.34E-34** |
| CCSp CE | All (28,026) | 3,608 | 3,281 | 0.161 | 0.009 | **2.43E-70** | 0.163 | 0.011 | **1.05E-49** |
| TOAST CE | Non-AF (3,869) | 3,333 | 3,074 | 0.171 | 0.013 | **3.17E-37** | 0.172 | 0.015 | **3.22E-29** |
| TOAST CE | All (28,026) | 3,333 | 3,074 | 0.149 | 0.009 | **3.00E-56** | 0.146 | 0.011 | **4.43E-41** |
| *Results after standardizing PRS to a z-score* |  |  |  |  |  |  |  |  |  |
| CCSc CE | Non-AF (3,869) | 3,000 | 2,725 | 0.365 | 0.027 | **1.59E-42** | 0.425 | 0.035 | **1.40E-34** |
| CCSc CE | All (28,026) | 3,000 | 2,725 | 0.329 | 0.019 | **1.01E-65** | 0.337 | 0.023 | **1.45E-48** |
| CCSp CE | Non-AF (3,869) | 3,608 | 3,281 | 0.348 | 0.025 | **6.98E-43** | 0.397 | 0.033 | **8.34E-34** |
| CCSp CE | All (28,026) | 3,608 | 3,281 | 0.315 | 0.018 | **2.43E-70** | 0.318 | 0.021 | **1.05E-49** |
| TOAST CE | Non-AF (3,869) | 3,333 | 3,074 | 0.334 | 0.026 | **3.17E-37** | 0.335 | 0.030 | **3.22E-29** |
| TOAST CE | All (28,026) | 3,333 | 3,074 | 0.291 | 0.018 | **3.00E-56** | 0.284 | 0.021 | **4.43E-41** |

| Case definition | Control definition (N) | Cases |  | Logistic regression, adjusted for PCs and sex |  |  | Logistic regression, adjusted for PCs, sex, and clinical covariates |  |  |
| --- | --- | --- | --- | --- | --- | --- | --- | --- | --- |
|  |  |  |  | Beta | SE | P-value | Beta | SE | P-value |
| CCSc SAO | Non-AF (3,869) | 2,262 | 2,124 | 0.023 | 0.017 | 0.170 | 0.026 | 0.019 | 0.163 |
| CCSc SAO | All (28,026) | 2,262 | 2,124 | 0.002 | 0.012 | 0.842 | 0.006 | 0.013 | 0.660 |
| CCSp SAO | Non-AF (3,869) | 2,419 | 2,267 | 0.025 | 0.016 | 0.124 | 0.029 | 0.018 | 0.109 |
| CCSp SAO | All (28,026) | 2,419 | 2,267 | 0.003 | 0.012 | 0.787 | 0.007 | 0.013 | 0.602 |
| TOAST SAO | Non-AF (3,869) | 2,631 | 2,415 | 0.021 | 0.016 | 0.209 | 0.019 | 0.018 | 0.289 |
| TOAST SAO | All (28,026) | 2,631 | 2,415 | 0.001 | 0.011 | 0.902 | 0.003 | 0.013 | 0.826 |
| *Results after standardizing PRS to a z-score* |  |  |  |  |  |  |  |  |  |
| CCSc SAO | Non-AF (3,869) | 2,262 | 2,124 | 0.046 | 0.033 | 0.170 | 0.051 | 0.036 | 0.163 |
| CCSc SAO | All (28,026) | 2,262 | 2,124 | 0.005 | 0.023 | 0.842 | 0.012 | 0.026 | 0.660 |
| CCSp SAO | Non-AF (3,869) | 2,419 | 2,267 | 0.049 | 0.032 | 0.124 | 0.057 | 0.035 | 0.109 |
| CCSp SAO | All (28,026) | 2,419 | 2,267 | 0.006 | 0.023 | 0.787 | 0.013 | 0.025 | 0.602 |
| TOAST SAO | Non-AF (3,869) | 2,631 | 2,415 | 0.040 | 0.032 | 0.209 | 0.037 | 0.035 | 0.289 |
| TOAST SAO | All (28,026) | 2,631 | 2,415 | 0.003 | 0.022 | 0.902 | 0.005 | 0.025 | 0.826 |

| Case definition | Control definition (N) | Cases |  | Logistic regression, adjusted for PCs and sex |  |  | Logistic regression, adjusted for PCs, sex, and clinical covariates |  |  |
| --- | --- | --- | --- | --- | --- | --- | --- | --- | --- |
|  |  |  |  | Beta | SE | P-value | Beta | SE | P-value |
| CCSc UNDETER | Non-AF (3,869) | 4,574 | 4,169 | 0.036 | 0.013 | **0.004** | 0.031 | 0.014 | 0.022 |
| CCSc UNDETER | All (28,026) | 4,574 | 4,169 | 0.021 | 0.009 | 0.013 | 0.021 | 0.010 | 0.030 |
| CCSc INCUNC | Non-AF (3,869) | 2,280 | 2,093 | 0.046 | 0.016 | **0.003** | 0.045 | 0.017 | 0.010 |
| CCSc INCUNC | All (28,026) | 2,280 | 2,093 | 0.028 | 0.012 | 0.015 | 0.029 | 0.013 | 0.025 |
| CCSc CRYPTCE | Non-AF (3,869) | 2,294 | 2,076 | 0.030 | 0.016 | 0.051 | 0.026 | 0.017 | 0.124 |
| CCSc CRYPTCE | All (28,026) | 2,294 | 2,076 | 0.015 | 0.012 | 0.212 | 0.017 | 0.013 | 0.192 |
| CCSp Crypto | Non-AF (3,869) | 1,096 | 972 | 0.035 | 0.020 | 0.090 | 0.029 | 0.022 | 0.195 |
| CCSp Crypto | All (28,026) | 1,096 | 972 | 0.019 | 0.016 | 0.258 | 0.021 | 0.018 | 0.245 |
| TOAST UNDETER | Non-AF (3,869) | 3,479 | 3,216 | 0.033 | 0.013 | 0.015 | 0.028 | 0.014 | 0.055 |
| TOAST UNDETER | All (28,026) | 3,479 | 3,216 | 0.021 | 0.010 | 0.027 | 0.022 | 0.011 | 0.042 |
| *Results after standardizing PRS to a z-score* |  |  |  |  |  |  |  |  |  |
| CCSc UNDETER | Non-AF (3,869) | 4,574 | 4,169 | 0.071 | 0.025 | **0.004** | 0.061 | 0.027 | 0.022 |
| CCSc UNDETER | All (28,026) | 4,574 | 4,169 | 0.041 | 0.017 | 0.013 | 0.041 | 0.019 | 0.030 |
| CCSc INCUNC | Non-AF (3,869) | 2,280 | 2,093 | 0.090 | 0.030 | **0.003** | 0.088 | 0.034 | 0.010 |
| CCSc INCUNC | All (28,026) | 2,280 | 2,093 | 0.055 | 0.023 | 0.015 | 0.056 | 0.025 | 0.025 |
| CCSc CRYPTCE | Non-AF (3,869) | 2,294 | 2,076 | 0.059 | 0.030 | 0.051 | 0.051 | 0.033 | 0.124 |
| CCSc CRYPTCE | All (28,026) | 2,294 | 2,076 | 0.028 | 0.023 | 0.212 | 0.033 | 0.025 | 0.192 |
| CCSp Crypto | Non-AF (3,869) | 1,096 | 972 | 0.068 | 0.040 | 0.090 | 0.057 | 0.044 | 0.195 |
| CCSp Crypto | All (28,026) | 1,096 | 972 | 0.036 | 0.032 | 0.258 | 0.041 | 0.035 | 0.245 |
| TOAST UNDETER | Non-AF (3,869) | 3,479 | 3,216 | 0.064 | 0.026 | 0.015 | 0.054 | 0.028 | 0.055 |
| TOAST UNDETER | All (28,026) | 3,479 | 3,216 | 0.042 | 0.019 | 0.027 | 0.042 | 0.021 | 0.042 |

UNDETER, undetermined; INCUNC, incomplete and unclassified; CRYPTCE, cryptogenic and CE minor; Crypto, cryptogenic

**Supplementary Table 6.** Sensitivity analysis for the atrial fibrillation polygenic risk score. As a sensitivity analysis for the polygenic risk score (PRS), we constructed 3 additional PRSs, including SNPs +/-25kb, +/-50kb, and +/-100kb from the SNPs included in the original score. All scores remain highly significant when tested for association with cardioembolic stroke (using a logistic regression model). P-values after additionally adjusting for clinical covariates are also shown. Clinical covariates: age, cardiovascular disease, type 2 diabetes status, smoking status, and hypertension.

| PRS SNPs | Filters | Total SNPs | PRS p-value |  |
| --- | --- | --- | --- | --- |
|  |  |  | *Adjusted for PCs, sex* | *Adjusted for PCs, sex, clinical covariates* |
| Original SNPs | MAF > 1%<br>Info > 0.8 | 975 | $1.01 \times 10^{-65}$ | $1.44 \times 10^{-48}$ |
| Original SNPs +/- 25kb | MAF > 1%<br>Info > 0.8 | 146,631 | $9.13 \times 10^{-50}$ | $1.32 \times 10^{-37}$ |
| Original SNPs +/- 50kb | MAF > 1%<br>Info > 0.8 | 258,870 | $5.76 \times 10^{-48}$ | $1.40 \times 10^{-36}$ |
| Original SNPs +/- 100kb | MAF > 1%<br>Info > 0.8 | 462,146 | $4.47 \times 10^{-44}$ | $1.77 \times 10^{-32}$ |

**Supplementary Table 7.** Clinical covariates available in the SiGN data. We adjusted our analyses of a polygenic risk score for a series of clinical covariates that are associated with atrial fibrillation. Summary-statistics on these covariates are shown below for those samples classified as (a) cardioembolic stroke or (b) undetermined stroke. The number of samples with missing data are provided in parentheses where relevant.

*Cardioembolic*
| Phenotype | CCS Causative | CCS Phenotypic | TOAST |
| --- | --- | --- | --- |
| Female | 1,588 | 1,859 | 1,618 |
| Male | 1,247 | 1,541 | 1,520 |
| Age: mean (sd) | 74.7 (12.4) | 74.5 (12.3) | 71.0 (15.1) |
| Hypertensive (missing) | 2,195 (18) | 2,665 (21) | 2,272 (16) |
| Diabetes mellitus (missing) | 763 (26) | 950 (29) | 799 (8) |
| CAD (missing) | 989 (64) | 1206 (83) | 911 (119) |
| Smoking |  |  |  |
| Current | 379 | 468 | 513 |
| Former | 694 | 865 | 776 |
| Never | 1,737 | 2,055 | 1,905 |
| Total | 3,000 | 3,608 | 3,333 |

*Undetermined*
| Phenotype | CCS Causative | CCS Causative | CCS Causative | CCS Phenotypic | TOAST |
| --- | --- | --- | --- | --- | --- |
| Female | 1,880 | 1,024 | 856 | 420 | 1,445 |
| Male | 2,151 | 1,014 | 1,137 | 543 | 1,635 |
| Age: mean (sd) | 63.9 (15.4) | 67.7 (13.9) | 69.0 (15.9) | 58.9 (15.7) | 63.7 (16.1) |
| Hypertensive (missing) | 2,833 (23) | 1,512 (14) | 1,321 (9) | 612 (3) | 2,110 (29) |
| Diabetes mellitus (missing) | 958 (26) | 513 (14) | 445 (12) | 202 (4) | 708 (25) |
| CAD (missing) | 739 (169) | 421 (86) | 318 (83) | 115 (46) | 573 (100) |
| Smoking |  |  |  |  |  |
| Current | 1,090 | 582 | 508 | 239 | 813 |
| Former | 1,050 | 516 | 534 | 235 | 772 |
| Never | 2,202 | 1,081 | 1,121 | 548 | 1,711 |
| Total | 4,574 | 2,280 | 2,294 | 1,096 | 3,479 |

**Supplementary Table 8:**
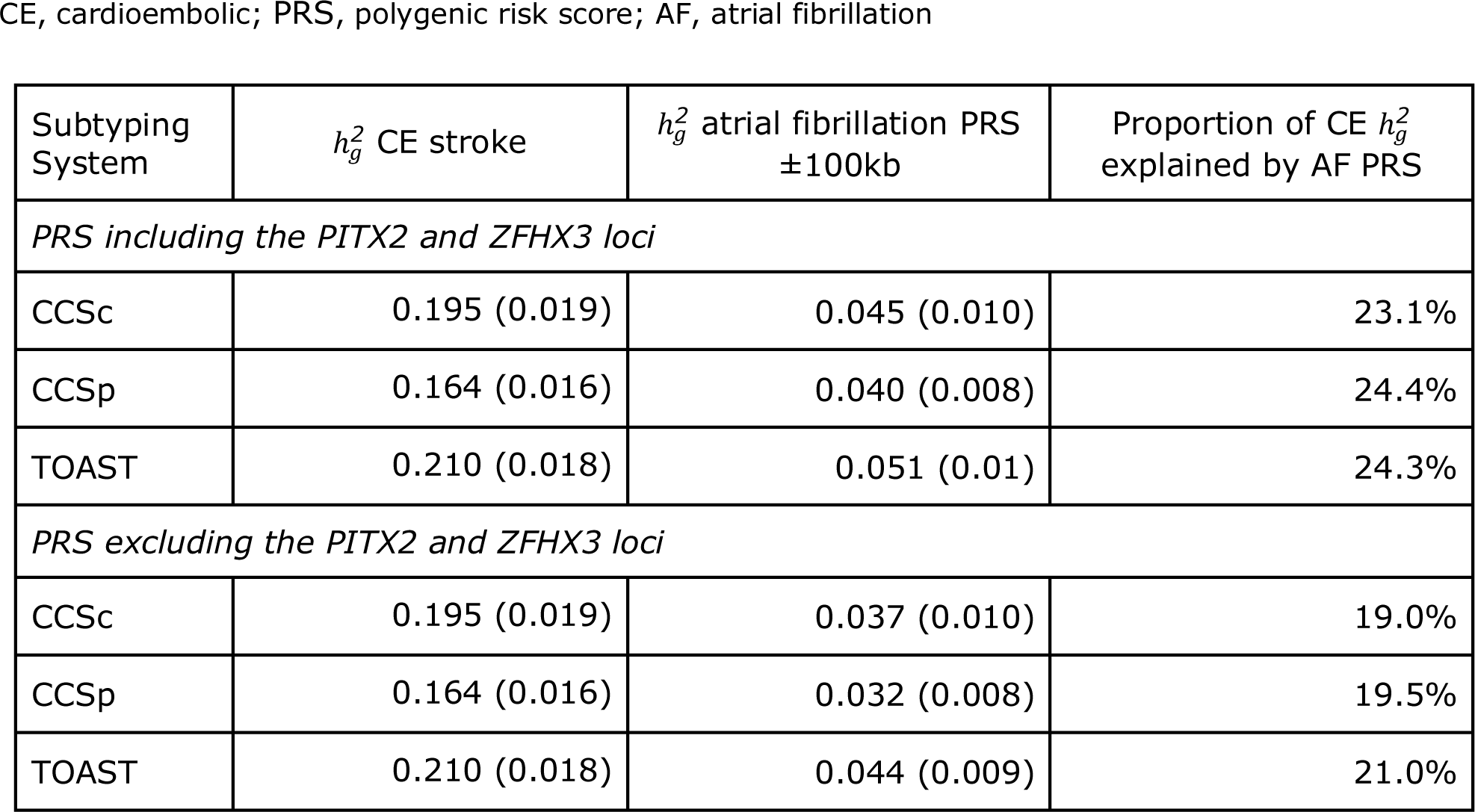
Variance explained by the atrial fibrillation polygenic risk score in cardioembolic stroke. To determine the variance explained by the atrial fibrillation polygenic risk score (PRS) in cardioembolic stroke, we constructed a model in BOLT-LMM that consisted of two variance components: (1) a variance component made up of SNPs for the genetic relationship matrix, and (2) a variance component made up of SNPs from the PRS. After computing the estimated variance explained for each component in BOLT-LMM, we converted the estimate to the liability score. Below is variance explained for each of the cardioembolic stroke phenotypes as determined by the three subtyping systems available in SiGN: CCS Causative, CCS Phenotypic, and TOAST. Standard errors of each estimate appear in parentheses. Explained variance is shown for a PRS including the *PITX2* (chromosome 4) and *ZFHX3* (chromosome 16) loci, as well as excluding ±2Mb around these loci (see https://github.com/UMCUGenetics/Afib-Stroke-Overlap for lists of SNPs that fall in these regions). Because a large number of SNPs is needed to construct a variance component to calculate variance explained, we performed the calculation using the atrial fibrillation PRS including SNPs ±100kb from the original PRS SNPs, and then pruning SNPs a linkage disequilibrium of 0.2.

| Subtyping System | $h_g^2$ CE stroke | $h_g^2$ atrial fibrillation PRS $\pm 100\text{kb}$ | Proportion of CE $h_g^2$ explained by AF PRS |
| --- | --- | --- | --- |
| *PRS including the PITX2 and ZFHX3 loci* |  |  |  |
| CCSc | 0.195 (0.019) | 0.045 (0.010) | 23.1% |
| CCSp | 0.164 (0.016) | 0.040 (0.008) | 24.4% |
| TOAST | 0.210 (0.018) | 0.051 (0.01) | 24.3% |
| *PRS excluding the PITX2 and ZFHX3 loci* |  |  |  |
| CCSc | 0.195 (0.019) | 0.037 (0.010) | 19.0% |
| CCSp | 0.164 (0.016) | 0.032 (0.008) | 19.5% |
| TOAST | 0.210 (0.018) | 0.044 (0.009) | 21.0% |

## Supplementary Methods

### GitHub repository and data availability

#### 1 GitHub repository and additional supporting data

*Relevant code for the analyses performed in this paper can be found here: https://github.com/saralpulit/Afib-Stroke-Overlap*.

This repository primarily consists of:

Call to BOLT-LMM to run GWAS

Call to GCTA and BOLT-LMM to calculate heritability

Call to PLINK^3,4^ to calculate the polygenic risk score (PRS)

An R script for converting observed heritability in BOLT-LMM to the liability scale (see below)

A script in R to check association between the PRS and various phenotypes

A call to PLINK^3,4^ to calculate a GRM to run GCTA

Sample identifiers for those individuals analyzed in this paper

SNP identifiers and weights for those markers included in the construction of the polygenic risk score

A complete README accompanies the GitHub repository.

#### 2 Sample and SNP identifiers used in these analyses

A file containing:

the dbGaP sample identifiers

the cohort the sample is drawn from

the continental group the sample is in (as determined in the first SiGN GWAS effort^5^)

a list of quality control-passing SNPs used in the initial GWAS is available on this paper’s GitHub repository.

#### 3 Downloadable summary-level genome-wide association study data

The summary-level data from the original SiGN GWAS has been made publicly available through the Cerebrovascular Disease Knowledge Portal, which can be accessed here: http://www.cerebrovascularportal.org/

These summary-level results are available for cardioembolic stroke (CE), large artery atherosclerosis (LAA), small artery occlusion (SAO), and undetermined (UNDETER) stroke, for three different subtyping systems (TOAST, CCS Causative, CCS Phenotypic).

The summary-level results for the atrial fibrillation genome-wide association studies (performed in broadly-defined or strictly-defined cases versus all controls) are available here:

Broadly-defined atrial fibrillation cases vs. all referents: https://doi.org/10.5281/zenodo.1035871

Strictly-defined atrial fibrillation cases vs. all referents: https://doi.org/10.5281/zenodo.1035873

### The Stroke Genetics Network (SiGN) and genome-wide association study of ischemic stroke subtypes

The full list of cohorts that are included in the SiGN genome-wide association study can be found in the Supplementary Material of “Loci associated with ischaemic stroke and its subtypes (SiGN): a genome-wide association study,”^5^ which can be accessed here: https://paperpile.com/shared/nvNXQf.

SiGN is comprised of several case cohorts with pre-existing genotyping data. Newly-collected cases, as well as a small number of matched referents, were genotyped on the Illumina 5M array^6^. The majority of referents included were drawn from publicly-available genotyping data.

#### 1 Referent (control) datasets

Referent datasets downloaded from the Database of Genotypes and Phenotypes (dbGaP) are:

|  | dbGAP accession # |
| --- | --- |
| Genetics Resource with the Health and Retirement Study | phs000428.v2.p2 |
| Whole Genome Association Study of Visceral Adiposity in the HABC study | phs000169.v1.p1 |

#### 2 Case datasets

A large number of cases and a small number of controls (from Belgium and Poland) were genotyped at the initiation of the SiGN GWAS. These data have been uploaded to dbGaP and are available here:

The National Institute of Neurological Disorders and Stroke (NINDS) Stroke Genetics Network (SiGN) (phs000615.v1.p1)

#### 3 Phenotyping in SiGN

There are three primary subtype definitions of ischemic stroke: cardioembolic stroke, large artery atherosclerotic stroke, and small artery occlusion. The SiGN consortium used the CCS system to attempt to assign each case to one of these three categories. Additionally, ∼74% of cases were also classified using the Trial of Org 10 172 in Acute Stroke Treatment (TOAST)^7,8^ system, which classifies stroke cases based on clinical decision-making and clinically-ascertained information. The CCS and TOAST subtyping systems yield moderately-to-strongly correlated phenotyping results (**Supplementary Figure 5**)^9^. Use of these traits in a GWAS setting also yields concordant association results, as previously shown ^6^. These subtypes are similarly defined in CCS and TOAST, though determined differently across the two subtyping systems.

In addition to the three primary subtypes, both the CCS and TOAST classification systems generate two additional subtypes: “undetermined” and “other.” The “other” classification was small in sample size (N_cases_ = 595, 719 and 374 in CCS Causative, CCS Phenotypic and TOAST, respectively), and was therefore not included in the original SiGN GWAS and was not tested here^6^. The “undetermined” classification, though named the same in CCS and TOAST, is defined differently across the two subtyping systems^8,10^. In TOAST, patients with conflicting subtype classifications are placed in the undetermined category^6,8^. In contrast, the CCS undetermined classification includes patients with cryptogenic embolism, other cryptogenic cases, patients with an incomplete evaluation, or samples with competing subtypes^10^.

#### 4 Brief summary of data quality control in SiGN

SiGN samples represent three continental populations (European-ancestry; African-ancestry; and non-European ancestry and non-African ancestry samples, primarily of admixed ancestry from Latin American populations, labelled ‘Hispanic’). In total, the study contains 13 case-referent analysis groups: 10 of European ancestry, two of African ancestry, and one Hispanic^6^.

For quality control (QC) and downstream association testing, cases and referents were matched by genotyping array and PCA-determined ancestry. European-ancestry samples were imputed with IMPUTE2^11^ using a reference panel built from whole-genome sequence data collected by the 1000 Genomes Project (Phase 1)^12^ and the Genome of the Netherlands^13^ project; African-ancestry and Hispanic samples were imputed with the 1000 Genomes Project data only.^12^ Due to data-sharing restrictions regarding the referents used for the Hispanic set of samples, only the European- and African-ancestry samples were analyzed here, totaling 13,390 cases and 28,026 referents distributed across 12 case-control analysis groups.

Before performing genome-wide association testing, for those SNPs that were genotyped in a subset of the SiGN study strata but imputed in others, we compared the frequency of the SNP across the various strata. We removed any SNP with a frequency difference > 15% within ancestral group or >50% across ancestral groups comparing imputed and genotyped data, likely induced by sequencing errors in the imputation reference panel(s).

### Constructing a genetic relationship matrix for genome-wide association testing in BOLT-LMM

To construct the genetic relationship matrix (GRM) implemented in BOLT-LMM, we used SNPs that were (i) common (MAF > 5%), (ii) with missingness < 5%, (iii) linkage disequilibrium (LD) pruned at an r^2^ threshold of 0.2, (iv) on the autosomal chromosomes only, (v) and not in stratified areas of the genome (i.e., not in the major histocompatibility complex (MHC), the inversions on chromosomes 8 and 17, or in the lactase (*LCT*) locus on chromosome 2). After association testing, we additionally removed SNPs with imputation quality (info score) < 0.8, due to excess inflation of the test statistic in those SNPs (**Supplementary Figure 1**).

### Running a genome-wide association study using BOLT-LMM

We implemented a linear mixed model to perform association testing using BOLT-LMM.^14^ Linear mixed models can account for structure in the data, such as that due to (familial or cryptic) relatedness and population structure, while improving power for discovery.^15–17^ Due to extensive structure in the SiGN data,^6^ induced by both study design and population ancestry, we adjusted the BOLT-LMM model for the top ten principal components (PCs) and sex, in addition to the genetic relationship matrix used as a random effect in the linear mixed model.^14^ We calculated PCs in EIGENSTRAT^18^ using a similar set of SNPs to that used in the genetic relationship matrix but using a missingness threshold of 0.1%. To construct the GRM, we first identified the set of SNPs with imputation quality > 0.8 and MAF > 1%. More than 5.5M SNPs passed these QC criteria, so we randomly selected 20% of the data (∼1.1M SNPs) for computational efficiency in calculating the GRM. We also identified SNPs outside the MHC and *LCT* regions, outside the inversions on chromosomes 8 and 17, and LD pruned (r^2^ = 0.2). These filtering steps resulted in ∼250,000 SNPs available for the GRM. We used Plink 1.9^3,4^ to convert imputed dosages to best-guess genotypes and then compute the GRM.

### SNP-based heritability calculations in GCTA and BOLT-LMM

We used the GRM from our GWAS analyses (described in the section above) to estimate heritability. We adjusted all heritability analyses for 10 PCs and sex. To test the robustness of our heritability estimates, we calculated three additional GRMs to re-estimate heritability, and additionally estimated heritability using a second software (GCTA^2^).

To check the robustness of the heritability calculations to the SNPs included in the GRM, we calculated heritability using the GRM described above, as well as three additional GRMs: (i) using the ∼1.1M SNPs with imputation quality > 0.8 and MAF > 1% (and without LD pruning); (ii) using the SNPs that were genotyped across all study strata (∼155,000 SNPs); and (iii) the set of genotyped SNPs with the MHC, *LCT* locus, inversions on chromosomes 8 and 17 removed, and LD pruned at r^2^ = 0.2.

Additionally, we computed heritability in GCTA^2^ using the same GRMs and assuming a trait prevalence of 1%. We compared the results to the BOLT-based 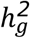 estimates (**Supplementary Table 3** and **Supplementary Figures 2-3**). As genome-wide heritability estimates need a large number of SNPs to be accurate, we report in the paper all estimates using a GRM containing imputed, pruned SNPs. Estimates resulting from all GRMs are presented here, in the **Supplementary Information**.

To test the effect of changing the GRM (referred to by the --bfile and ‘modelSNPs’ option in BOLT-LMM), we selected SNPs for the GRM in four ways:

(1) Genotyped SNPs only (minor allele frequency > 1%) (115,553 SNPs total)

(2) Genotyped SNPs, pruned at a linkage disequilibrium threshold (r2 threshold) of 0.2, and removing the MHC, *LCT* locus, and two chromosomal inversions. (60,432 SNPs total)

(3) Imputed SNPs (minor allele frequency > 1% and imputation info > 0.8) converted to best-guess genotypes. (1,128,985 SNPs total)

(4) Imputed SNPs (minor allele frequency > 1% and imputation info > 0.8); pruned at a linkage disequilibrium threshold (r2 threshold) of 0.2; removing the MHC, *LCT* locus, and two chromosomal inversions; and converted to best-guess genotypes. (250,209 SNPs total)

The GRM in (4) is the GRM used for all heritability results presented in the main manuscript.

As calculating GRMs in GCTA can be extremely computationally intensive, we calculated the GRMs using PLINK 1.9 and then used those GRMs to estimate heritability. A script that shows how to do this is included in the GitHub repository noted above.

The genomic locations (hg19) for excluded markers are as follows:

BOLT-LMM produces heritability estimates on the observed scale. To convert to the liability scale (i.e., the scale on which GCTA produces heritability estimates) we performed a conversion in R. Running the conversion requires knowing the trait prevalence, total cases analyzed, total controls analyzed, and the heritability on the observed scale. This code snippet is available in the accompanying GitHub repository for this paper.

### Quality control in genome-wide data for correlation calculations

We used summary-level data from the latest Atrial Fibrillation Genetics (AFGen) Consortium meta-analysis of atrial fibrillation^1^ to calculate a z-score for each SNP in that GWAS. Additionally, we calculated a z-score for each SNP in a GWAS of each stroke subtype in SiGN as well as in the GWAS of atrial fibrillation we performed in the SiGN data. Finally, as a null comparator, we downloaded SNP z-scores from a GWAS of educational attainment^19^available through LDHub (http://ldsc.broadinstitute.org/, accessed 11-1-2017). We aligned z-score signs based on the risk allele reported in each study. SNPs with an allele frequency difference >5% between AFGen and SiGN (all stroke analysis) were removed from the AFGen data (25,784 SNPs); similarly, SNPs with an allele frequency difference >5% between the educational attainment GWAS and SiGN (all stroke) were also removed (27,866 SNPs). Finally, we calculated Pearson’s r between z-scores from two traits to evaluate correlation.

### Constructing an atrial fibrillation polygenic risk score

To construct an atrial fibrillation polygenic risk score (PRS), we used SNPs from a previously-derived atrial fibrillation PRS.^20^ Briefly, the PRS was derived using results from a recent GWAS of atrial fibrillation, comprised of 17,931 cases and 115,142 referents^1^ and testing various sets of SNPs based on their p-value from that GWAS (varying from p < 5 × 10^−8^ to p < 0.001) and using varied linkage disequilibrium thresholds (0.1 - 0.9).^20^ These sets of SNPs were used to generate various PRSs, which were then independently tested for association to atrial fibrillation in an independent sample from the UK Biobank; the best-performing PRS (defined as the PRS with the lowest Akaike’s Information Criterion) comprised 1,168 SNPs with p < 1 × 10^−4^ in the atrial fibrillation GWAS and LD pruned at an r^2^ threshold of 0.5.^20^

Of these 1,168 SNPs, we identified 934 SNPs in the SiGN dataset with imputation info

> 0.8 and MAF > 1%. We used these 934 SNPs to construct the atrial fibrillation PRS in the SiGN dataset by weighting the imputed number of risk-increasing alleles carried by an individual at a given SNP (i.e., 0-2 risk-increasing alleles) and then weighting the dosage by the effect of the allele, as determined by the most recent GWAS.^1^ We computed the final PRS for each individual by summing across all of the weighted genotypes and performed association testing in R.

We calculated the odds ratio of the PRS for an increase of one standard deviation in the score by first converting the PRS per individual to a z-score, where:

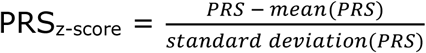

We then recalculated the association between PRS_z-score_ and the phenotype, and converted the resulting regression coefficients (i.e., betas) of the PRS to odds ratios.

To ensure that our analyses of the PRS were robust to ancestral heterogeneity, we additionally tested the PRS in the subset of European-ancestry samples only (the data were essentially identical to our finding in the complete sample and are therefore not provided).

## Supplementary Results

### Including age as a covariate in the GWAS of atrial fibrillation

To check for the effects of age on our initial GWAS findings, we ran a GWAS of atrial fibrillation including age as a covariate. Controls without age information were dropped from this analysis. Given the structure of the SiGN dataset -- which includes groups of cases and controls that have been carefully matched on genotyping array and ancestry -- we also dropped the cases for which their matched controls were missing age information.

Our age-adjusted analysis included 2,487 atrial fibrillation cases and 22,072 controls. We performed the GWAS in BOLT-LMM, adjusting for 10 PCs, sex and age. We then checked the correlation between the SNP effects (betas) from the GWAS unadjusted for age and the SNP effects from the GWAS adjusted for age. Correlation was strong (r = 0.83).

## Appendix I

Members of the Atrial Fibrillation Genetics (AFGen) Consortium

Please note that the AFGen Consortium participants evolve over time. Further information on the AFGen Consortium can be found at www.afgen.org.

## Appendix II

Members of the International Stroke Genetics Consortium (ISGC)

Please note that ISGC participants evolve over time. Further information on the ISGC can be found at http://www.strokegenetics.org/.

Author Contributions
All named authors have contributed meaningfully to the present study. Specific contributions for each author are described below.
**S.L. Pulit:** conception of research design, data analysis, drafting of manuscript, critical revision of manuscript
**L.C. Weng:** data analysis, critical revision of manuscript
**P.F. McArdle:** data acquisition, data analysis, critical revision of manuscript
**L. Trinquart:** data acquisition, critical revision of manuscript
**S.H. Choi:** data acquisition, critical revision of manuscript
**B.D. Mitchell:** data acquisition, study supervision, critical revision of manuscript
**J. Rosand:** data acquisition, study supervision, critical revision of manuscript
**P.I.W. de Bakker:** study supervision, critical revision of manuscript
**E.J. Benjamin:** data acquisition, study supervision, critical revision of manuscript
**P.T. Ellinor:** data acquisition, study supervision, critical revision of manuscript
**S.J. Kittner:** data acquisition, study supervision, critical revision of manuscript
**S.A. Lubitz:** conception of research design, study supervision, drafting of manuscript, critical revision of manuscript
**C.D. Anderson:** conception of research design, study supervision, drafting of manuscript, critical revision of manuscript

